# A paradigm shift of SERPINA3N in neurobehavioral development and brain injury

**DOI:** 10.1101/2025.09.09.675167

**Authors:** Meina Zhu, Bhabotosh Barman, Chengda Guo, Rohini Murali, Jane Joseph, Fuzheng Guo

## Abstract

Murine serine protease inhibitor clade A member 3N (SERPINA3N) and its human ortholog SERPINA3 are dysregulated in neurological disorders. SERPINA3N has been proposed as a protective factor, enhancing neurobehavioral development under homeostasis and mitigating neuronal/glial and vascular damage under neurological conditions. Here, we employed powerful, non-invasive genetic tools to revisit the concept of SERPINA3N in brain development and injury. Brain-specific SERPINA3N overexpression neither alters neuronal or glial development nor improves cognitive ability under homeostasis. In stark contrast, SERPINA3N drives a pro-inflammatory response to brain injury and exacerbates blood-brain barrier dysfunction. Its overexpression promotes neurodegeneration through apoptosis-mediated neuronal loss and disrupts oligodendroglial differentiation and myelination following neonatal brain injury. Collectively, our findings challenge the prevailing paradigm of SERPINA3N in neurodevelopment and injury, revealing that while SERPINA3N is dispensable for neurobehavioral development, it aggravates neural injury and vascular damage under pathological conditions.

## Introduction

Murine serine protease inhibitor clade A member 3N (SERPINA3N, encoded by *Serpina3n*) and its human ortholog SERPINA3 ^1^ (α_1_-antichymotrypsin, encoded by *SERPINA3*) are acute-phase proteins secreted primarily by the liver in response to systemic inflammation ^2^. During normal brain development, the baseline expression of SERPINA3N/SERPINA3 is relatively low in neural precursor cells that give rise to neurons and glia ^3-5^. We recently used genetic tools to delete SERPINA3N (loss-of-function) from Olig2-expressing neural precursor cells and concluded that SERPINA3N is dispensable for brain development or animal behaviors ^6^. Nevertheless, a prior study reported that enforced expression of human SERPINA3 (gain-of-function) increased neurogenesis and enhanced cognitive function in mice ^3^. That study was built on the rationale that human SERPINA3 drives cortical expansion in mice because endogenous SERPINA3N, its murine ortholog, is expressed at undetectable levels in the embryonic mouse brain. If true, SERPINA3N gain-of-function should similarly expand neuronal populations and improve cognition in mice. However, to date, no studies have directly examined the impact of brain-specific SERPINA3N overexpression on mouse neurobehavioral development.

Beyond normal brain development, clinical interest began with the discovery that SERPINA3 is deposited to β-amyloid plaques of Alzheimer’s disease ^7^. Subsequent studies reported elevated SERPINA3 levels in the plasma and cerebrospinal fluid of patients with various neurological diseases and injuries (see review ^2^). Notably, elevated SERPINA3 levels often correlate with disease severity, progression, or poor outcomes ^8-13^, suggesting a potential pathogenic role. However, its role in regulating CNS pathophysiology remains controversial ^2^.

Many studies have suggested that SERPINA3N protects neurons and the blood-brain barrier from experimental damage and exerts anti-inflammatory effects during brain injury ^14-20^. Yet, these findings are incompatible with the clinical observation that elevated SERPINA3 level is associated with disease deterioration. In contrast to a protective role, a few studies suggest that SERPINA3N may cause neurodegeneration ^21^ and blood-brain barrier disruption^22^ *in vitro* and drive pro-inflammatory responses to epileptic injury *in vivo* ^23^. The reasons for the discrepant results remain elusive, yet experimental systems (such as *in vitro* vs. *ex vivo*, vs. *in vivo*) or SERPINA3N intervention approaches may play a major part. Therefore, it is imperative to use brain-specific genetic approaches to revisit the role of SERPINA3N in regulating neurodegeneration, neuroinflammation, and blood-brain barrier disruption, the shared pathophysiology of neurological diseases and injuries.

Here, we employed *Cre-loxP* genetic approaches to alter SERPINA3N specifically in brain neural cells and assess the *in vivo* effect of SERPINA3N expression on mouse neurobehavioral development and brain injury. We reported that SERPINA3N gain-of-function does not alter neurobehavioral development in mice, nor does it perturb blood-brain barrier function or elicit neuroinflammation during homeostasis. Under injury conditions, neural cell-derived SERPINA3N drives a pro-inflammatory response, promotes neuronal apoptosis and neurodegeneration, and disrupts oligodendroglial differentiation and myelination. Our findings challenge the popular paradigm of SERPINA3N in brain development and injury. We propose that neural cell-derived SERPINA3N plays a deleterious role aggravating brain pathophysiology.

## Results

### Generation of SERPINA3N overexpression transgenic mice

To study the biological outcomes of enforced SERPINA3N expression, we generated Serpina3n conditional overexpression (cOE) transgenic mice (referred to as *lsl-Serpina3n* (**Suppl Fig. 1**). To achieve brain-specific SERPINA3N expression, we crossed *lsl-Serpina3n* mice with *Nestin-Cre* mice ^3,24,25^. *Nestin-Cre* activity is observed in neural precursor cells (NPCs) beginning at embryonic day 10.5 (E10.5) and peaking at E12.5 ^25^. The resulting double transgenic mice *Nestin-Cre:lsl-Serpina3n* were viable and fertile and exhibited no abnormalities in gross behaviors during homeostasis.

### SERPINA3N expression does not affect neurodevelopment

We evaluated SERPINA3N expression in *Nestin-Cre:lsl-Serpina3n* mice (referred to as brain-specific Serpina3n cOE) at the weaning ages. Transgenic mice carrying *Nestin-Cre* only were used as Serpina3n Ctrl mice. Western blot assay showed a marked elevation of SERPINA3N in the brain and spinal cord but not in peripheral organs such as liver and thymus (**Fig. 1A**). Consistently, real time quantitative PCR (RT-qPCR) showed a >50-fold increase in *Serpina3n* mRNA levels in the brain (**Fig. 1B**). *Nestin-Cre*-mediated SREPINA3N expression is concomitant with EGFP expression (**Suppl Fig. 1A**) in Nestin^+^ NPCs and their neural progenies. Accordingly, many neural cells were positive for EGFP, such as SOX10^+^ oligodendrocytes and GFAP^+^ astrocytes in the postnatal brain of Serpina3n cOE mice (**Suppl Fig. 2A**). In contrast, EGFP is absent from IBA1^+^ microglia (**Suppl Fig. 2B**), which are of myeloid origin. Thus, SERPINA3N is specifically expressed in brain neural cells in Serpina3n cOE mice.

**Figure 1.**
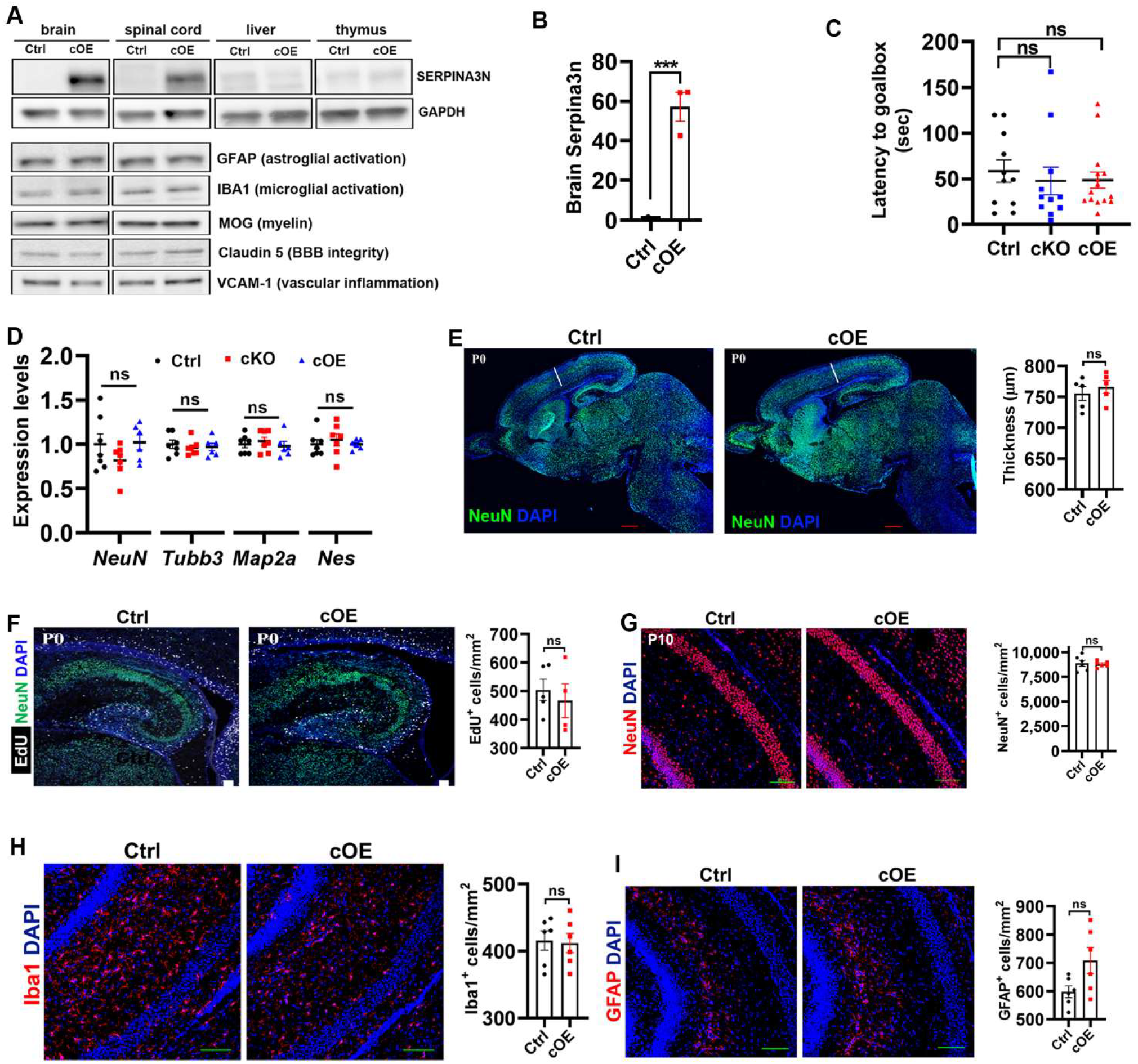
Serpina3n expression does not affect animal behavior, neurodevelopment, nor does it elicit glial cell activation during homeostasis. **A**, Western blot assays of SERPINA3N, GFAP, IBA1, MOG (myelin-oligodendrocyte glycoprotein), Claudin 5, and VCAM1 (vascular cell adhesion molecule 1) in brain, spinal cord, and/or peripheral organs of *Nestin-Cre:lsl-Serpina3n* (Serpina3n cOE) and *Nestin-Cre* Ctrl mice at P22. GAPDH used as an internal loading control. **B**, RT-qPCR quantification of brain Serpina3n mRNA at P22. Serpia3n levels were normalized to the housekeeping gene Hsp90. N=3, unpaired Student’s t test, t_(5)_ = 9.201, *P* = 0.0003. **C**, time latency to finding the goalbox during Barnes maze test at day 6 after training. cKO, *Nestin-Cre:Serpina3n*^fl/fl^. Two-month-old mice were used for testing. N=11 Ctrl, 11 cKO, 15 cOE. One-way ANOVA, *F*_(2,34)_ = 0.9219 *P* = 0.4075. **D**, RT-qPCR assays for neuronal markers NeuN, Tubb3, and Map2a and neural precursor cell NPC marker Nestin (Nes) in the brain of two-month-old mice. Two-month-old mice were used for testing. N=7 Ctrl, 7 cKO, 6 cOE. One-way ANOVA, *F*_(2,17)_ = 1.456 *P* = 0.2608 NeuN, *F*_(2,17)_ = 0.2855 *P* = 0.7552 Tubb3, *F*_(2,17)_ = 0.3609 *P* = 0.7-23 Map2a, *F*_(2,17)_ = 0.2990 *P* = 0.7454 Nes. **E**, representative images of neuronal marker NeuN and nuclear counterstaining DAPI and cortical thickness in the brain of newly born pups (P0). N=5, unpaired Student’s t test, t_(8)_ = 0.6902 *P* = 0.5096. **F**, representative images of NeuN and thymidine analog EdU (labeling proliferating cells, 2 hours pulse labeling) in the cortical and hippocampal areas of newly born pups. N=5 Ctrl, 4 cOE, unpaired Student’s t test, t_(7)_ = 0.5660 *P* = 0.5891. **G**, representative images of NeuN in the hippocampus of postnatal day 10 (P10) mice and quantification of NeuN^+^ neurons. N=6, unpaired Student’s t test, t_(10)_ = 0.2912 *P* = 0.7768. **H**, representative images and quantification of microglial marker IBA1in the hippocampus of P10 mice. N=6, unpaired Student’s t test, t_(10)_ = 0.1772 *P* = 0.8629. **I**, representative images and quantification of astrocyte marker GFAP in the hippocampus of P10 mice. N=6, unpaired Student’s t test, t_(10)_ = 2.189 *P* = 0.0534. Scale bars: E, 500µm, F, 50µm, G-I, 100µm.

Despite enforced SERPINA3N expression, no difference was observed in the brain levels of astroglial lineage GFAP (upregulation upon astroglial activation), IBA1 (upregulation upon microglial activation), tight junction protein Claudin5 (downregulation upon BBB disruption), VCAM1 (upregulation upon vascular inflammation), or MOG (downregulation upon oligodendroglial maldevelopment) (**Fig. 1A**). RT-qPCR assay did not reveal any difference in the mRNA levels of lineage-specific genes involving in oligodendrocyte differentiation and myelination (**Suppl Fig. 2C**), astrocyte maturation and activation (**Suppl Fig. 2D**), neuronal maturation and function (**Suppl Fig. 2E**), or microglial homeostasis and activation (**Suppl Fig. 2F**). The absence of microglial activation in Serpina3n cOE mice was confirmed by fluorescence immunostaining of microglial activation marker CD68 (**Suppl Fig. 2G**). Together, these data indicate that SERPINA3N expression does not perturb postnatal neural cell development, consistent with our recent conclusion that SERPINA3N is dispensable for brain development ^6^.

### SERPINA3N expression does not improve neurobehavioral development in adult mice

To determine if enforced SERPINA3N expression improves neurobehavioral development in mice as previously suggested ^3^, adult Serpina3n cOE and Ctrl mice were subjected to a battery of behavioral tests. Barnes maze test was used to evaluate animal’s cognitive function (learning and memory). Our results showed no significant difference in the latency to finding the goalbox during the probe session (**Fig. 1C**) between Serpina3n cOE and Ctrl mice while both groups exhibited a sharp learning curve during the 5-day training session (**Supple Fig. 3A**). These data suggest that SERPINA3N gain-of-function does not improve cognitive function in mice. Furthermore, both groups displayed similar motor performance on accelerating rotarod test (**Suppl Fig. 3B**). Next, we used open field test to assess the anxiety level and locomotor activity and found no perturbation in both metrics (**Suppl Fig. 3C**), which is further corroborated by elevated plus maze test (**Suppl Fig. 3D**). Finally, we generated *Nestin-Cre:Serpina3n*^fl/fl^ mice (Serpina3n cKO) to assess the effect of SERPINA3N loss-of-function on neurobehavioral development. Our data demonstrated that, like Serpina3n cOE mice, Serpina3n cKO mice exhibited no impaired ability in cognition, motor, and anxiety compared with Serpina3n Ctrl or cOE mice (**Fig. 1C, Suppl Fig. 3)**. Thus, SERPINA3N plays a dispensable role in neurobehavioral development.

### Normal brain hemostasis in adult mice of Serpina3n gain-of-function and loss-of-function

The unaltered neurobehaviors in Serpina3n cOE and cKO mice indicate normal brain cell development. We used RT-PCR to quantify lineage-specific genes indicative of cell development and activation and found no difference in the expression levels of neuronal markers NeuN, Tubb3 and Map2a and neural precursor marker NPC Nestin (**Fig. 1D**), oligodendroglial markers (Sox10, Mbp, Mobp, Myrf) (**Suppl Fig. 4A**), microglial homeostasis and activation markers (Tmem119, Iba1, Cd68) (**Suppl Fig. 4B**) or astroglial marker (Gfap, Adlh1l1) (**Suppl Fig. 4C**). These results suggest that brain homeostasis is properly maintained in adult Serpina3n cOE and cKO mice.

### SERPINA3N expression does not perturb neuronal and glial cell development

The unaltered brain homeostasis in adult mice suggests a normal trajectory of neuronal/glial cell development. Alternatively, neuronal/glial cell development may be temporally disrupted during early brain development and compensate to the normal level later in the adult. To test these two possibilities, we assessed neuronal/glial cell development in Serpina3n cOE mice during late embryonic and early postnatal ages. At postnatal day 0 (P0) when most, if not all, cortical neurons are already generated and migrated into their respective destinations, the cerebral cortical thickness of Serpina3n cOE mice was indistinguishable from that of littermate Ctrl mice (**Fig. 1E**). We did not observe cortical expansion or folding in Serpina3n cOE mice, a prominent cortical phenotype as previously reported ^3^. Cortical NPCs are actively proliferating in the ventricular zone and subventricular zone at E15.5 ^26^. However, at this developmental stage, we did not find difference in EdU^+^ proliferating cells in those zones of Serpina3n cOE brain compared with Ctrl brain (**Suppl Fig. 5A**). Abundant Nestin^+^ NPCs are proliferating in the hippocampus during late embryonic and early postnatal ^25^. No difference was observed in the density of EdU^+^ proliferating cells in the hippocampus of Serpina3n cOE neonates compared with that of Ctrl littermates (**Fig. 1F, Suppl Fig. 5B**). Consistently, the density of NeuN^+^ neurons was indistinguishable between Serpina3n cOE and littermate Ctrl mice during early postnatal development (**Fig. 1G, Suppl Fig. 6A**). Thus, SERPINA3N expression does not perturb neural precursor cell proliferation or neuronal population generation.

Glial cell maturation (oligodendroglia, astroglia, and microglia) occurs predominantly during early postnatal development in the murine brain. We used P10 mice to assess glial cell maturation and potential activation. Our histological analysis revealed that the density of Iba1^+^ microglia (**Fig. 1H, Suppl Fig. 6B**) and GFAP^+^ astroglia (**Fig. 1H**) was statistically similar in Serpina3n cOE mice to that in Serpina3n Ctrl mice. We did not observe any features of microglial/astroglial activation, such as morphological hypertrophy or process retraction, in Serpina3n cOE mice. The densities of PDGFRa^+^ oligodendrocyte progenitor cells (OPCs, **Suppl Fig. 6C**) and CC1^+^ differentiated oligodendrocytes (OLs, **Suppl Fig. 6D**) were comparable in the subcortical white matter tract between Serpina3n cOE and Ctrl mice. Consistently, oligodendroglial myelination, visualized by MBP histological staining, was indistinguishable between the two groups (**Suppl Fig. 6E**). Taken together, these findings suggest that SERPINA3N expression does not affect the normal trajectory of neuronal/glial cell development during homeostasis nor cause microglial and astroglial activation.

### Genetic expression of SERPINA3N does not elicit BBB dysfunction or neuroinflammation during homeostasis

A prior study reported that exogenously administered SERPINA3N resulted in BBB dysfunction in the homeostatic brain ^22^. Our genetic model provides an exquisite tool to test if SERPINA3N elicits BBB dysfunction in the homeostatic brain without any external CNS trauma. We found that the occupying area of the tight junction protein Claudin-5 ^27^ along Laminin B1-labeled blood vessels was statistically comparable between Serpina3 cOE and Ctrl mice (**Suppl Fig. 7A**), suggesting that neural cell-derived SERPINA3N does not affect the physical formation of the BBB. Macromolecules such as albumin are circulating within the blood stream and do not cross over the functional BBB into the brain parenchyma. Thus, we use albumin immunostaining on perfused brain sections (removing albumin in the bloodstream) to assess the potential leakage of the BBB. Our results showed little signal of Albumin in the brain parenchymal area beyond Laminin B^+^ vasculature (**Suppl Fig. 7B**). Importantly, albumin-occupying area was indistinguishable in the brain between Serpina3n cOE and Ctrl mice (**Suppl Fig. 7C**), indicating a normal BBB formation and function. The expression of proinflammatory cytokines (IL1β and TNFα) and chemokines (CCL2 and CXCL10) was comparable in the brain between Serpina3n cOE and Ctrl mice (**Suppl Fig. 8A-B**). Similarly, no difference was observed in the expression of microglial homeostasis marker Tmem119 and activation marker CD68 (**Suppl Fig. 8C**) or pan-reactive astrocyte makers GFAP and Vimentin (**Suppl Fig. 8D**). Taken together, these results indicate that SERPINA3N overexpression does not affect BBB function, nor does it cause microglial/astroglial activation under homeostasis.

### SERPINA3N is dysregulated after brain injury

Having established that SERPINA3N expression is dispensable for neurodevelopment and glial development under homeostasis, we next asked if SERPINA3N is an injury-induced factor modulating brain pathophysiology under diseased conditions. To this end, we evaluated the role of SERPINA3N in neonatal hypoxic-ischemic encephalopathy (HIE). HIE is a serious birth complication affecting full-term infants caused primarily by hypoxia (low oxygen) and ischemia (impaired cerebral blood flow) ^28^. Brain pathophysiology of HIE includes BBB disruption ^29^, inflammation and microglia/astroglial activation ^30^, neuronal apoptosis and neurodegeneration ^31^, and diffused white matter injury and hypomyelination ^32^, which are shared pathological features in many neurological diseases/injuries. We created the mouse model of HIE by permanently ligating the right common carotid artery (ischemia) of P6 neonatal mice followed by 45 min 6% oxygen exposure (hypoxia) ^33^.

Given that SERPINA3N is a secreted protein, we generated *Serpina3n-tdTom* reporter mice to study SERPINA3N dysregulation after brain injury. In our Serpina3n -tdTom mice, the tdTomato coding sequence is knocked in the endogenous Serpina3n locus, replacing the STOP codon of murine *Serpina3n gene* (**Suppl Fig. 9A**). Time-course analysis showed that tdTom was sharply induced by day 1 post-hypoxic/ischemic injury (D1 post-H/I) primarily in the hippocampus and subcortical white matter tract, peaked at D3, and downregulated by D7 post-H/I (**Suppl Fig. 9B-C**). These findings clearly demonstrate that SERPINA3N is an injury-induced molecule and suggest that it may play a crucial role in regulating brain pathophysiology after H/I injury.

### SERPINA3N aggravates BBB dysfunction under pathological conditions

It remains elusive if the injury-induced SERPINA3N is a protective or deleterious factor for BBB function after brain injury although it is dispensable for BBB function during homeostasis. We observed a severe disruption in BBB function at D3 post-H/I injury as demonstrated by decreased Claudin-5 expression in the vasculature (**Fig. 2A**) and increased Albumin signal in the brain parenchyma (**Fig. 2D**), consistent with BBB disruption in HIE patients. Contrary to previous reports ^16,19,20^, genetic overexpression of SERPINA3N by neural cells significantly diminished the expression of tight junction protein Claudin-5 and concomitantly augmented the leakage of albumin into the brain parenchyma of H/I-injured Serpina3n cOE mice compared with Ctrl mice (**Fig. 2B, C, E**). These results establish that SERPINA3N aggravates the disruption of BBB integrity during brain injury.

**Figure 2.**
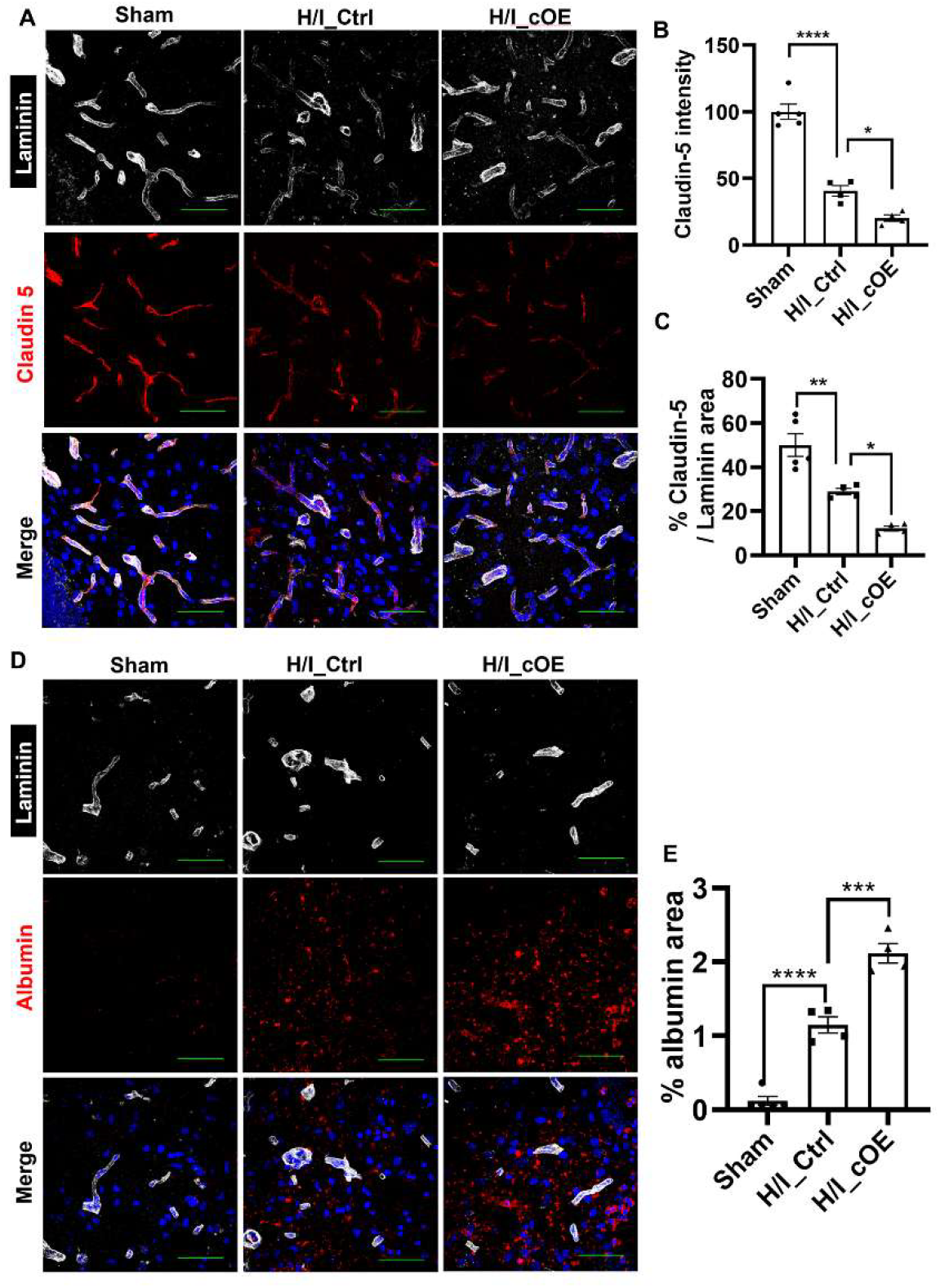
SERPINA3N expression aggravates BBB dysfunction under injury conditions. **A**, fluorescence immunostaining and showing alterations of tight junction protein Claudin 5 within Laminin-labeled blood vessels in the hippocampus at D3 post-H/I injury. Scale bars=50µm. **B**, Claudin 5 integrated intensity in the hippocampus at D3 post-H/I injury. N=5 sham, 4 H/I_Ctrl, 4 H/I_cOE. One-way ANOVA followed by Tukey’s multi-comparison test. *F*_(2,10)_ = 86.81 *P* < 0.0001, H/I_Ctrl vs. sham *P* < 0.0001, H/I_cOE vs. H/I_Ctrl *P* = 0.0309. **C**, percentage of Claudin 5-occupying area among laminin area. N=5 sham, 4 H/I_Ctrl, 4 H/I_cOE. One-way ANOVA followed by Tukey’s multi-comparison test. *F*_(2,10)_ = 28.33 *P* < 0.0001, H/I_Ctrl vs. sham *P* = 0.0052, H/I_cOE vs. H/I_Ctrl *P* = 0.0250. **D**, fluorescence immunostaining showing Albumin leakage into hippocampal parenchyma beyond blood vessels labelled by Laminin at D3 post-H/I injury. Scale bars=50µm. **E**, quantification of percentage of albumin in the parenchymal area. N=5 sham, 4 H/I_Ctrl, 4 H/I_cOE. One-way ANOVA followed by Tukey’s multi-comparison test. *F*_(2,10)_ =104.1 *P* < 0.0001, H/I_Ctrl vs. sham *P* < 0.0001, H/I_cOE vs. H/I_Ctrl *P* = 0.0002.

### SERPINA3N drives a pro-inflammatory response to brain injury and promotes microglial and astroglial activation

Neuroinflammation is a key pathophysiological feature of not only HIE but also other neurological diseases and injuries. Interleukin 1β (IL1β) and tumor necrosis factor α (TNFα) are potent pro-inflammatory cytokines that induce and perpetuate brain injury in HIE ^34-36^. We found that H/I injury caused >2-fold increase in IL1β expression and 100% increase in TNFα expression in the brain (**Fig. 3A**). SERPINA3N expression further increased the levels of IL1β and TNFα in the brain of H/I-injured Serpina3n cOE mice by 46% and 340%, respectively, compared with those of H/I-injured Serpina3n Ctrl mice (**Fig. 3A**). Chemokines CCL2 and CXCL10 increase the permeability of the BBB and contribute to the inflammatory cascade that causes brain damage ^37,38^. In line with the deleterious effect, the brain levels of CCL2 and CXCL10 were increased at D3 H/I injury (**Fig. 3B**). SERPINA3N gain-of-function dramatically increased the expression of CCL2 and CXCL10 in Serpina3n cOE mice compared with Ctrl mice during H/I injury (**Fig. 3B**). Together, these findings suggest that SERPINA3N drives a pro-inflammatory response to brain H/I injury.

**Figure 3.**
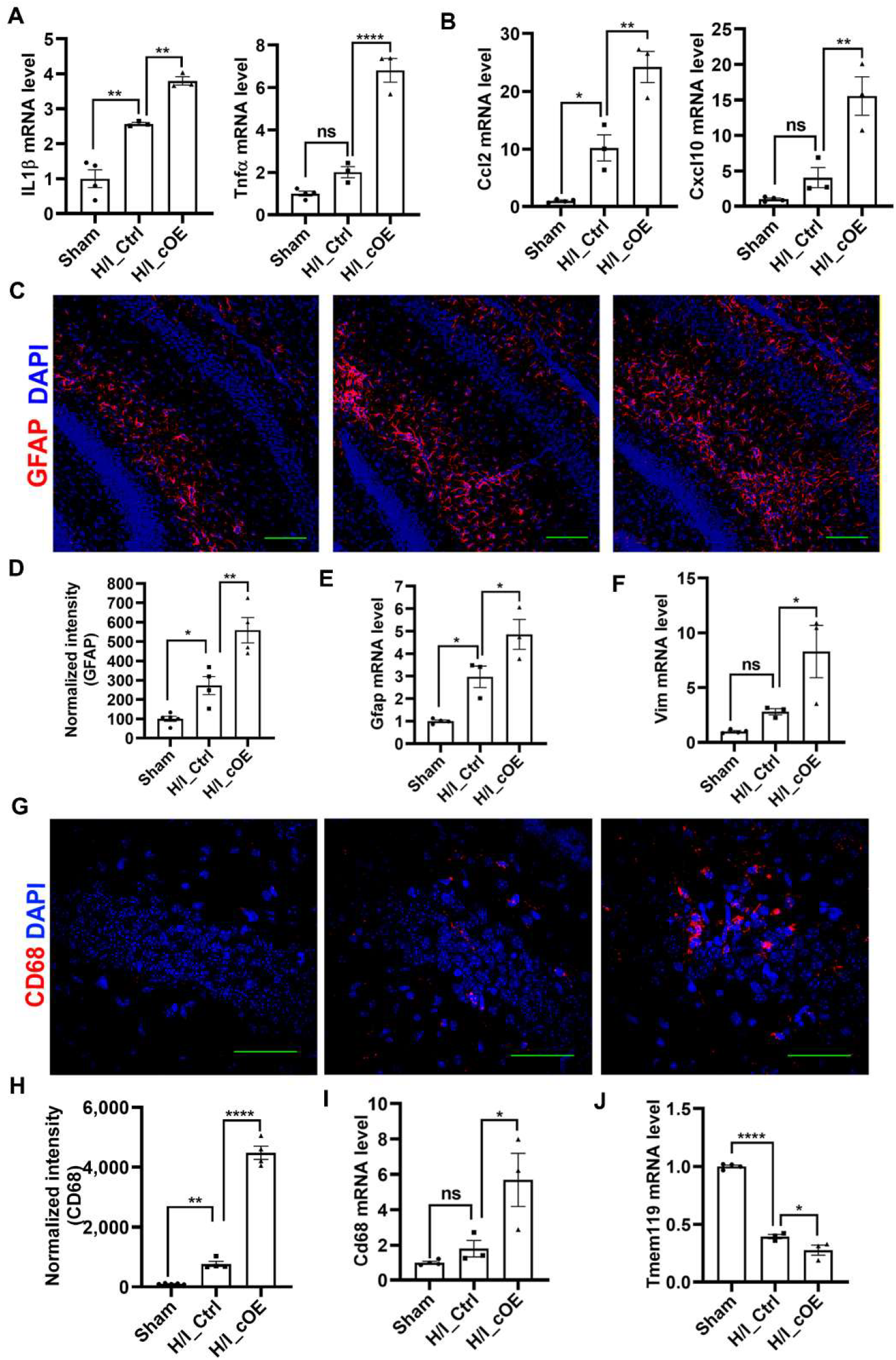
SERPINA3N regulates glial activation and neuroinflammation during H/I injury. **A**, RT-qPCR assays for brain expression of canonical pro-inflammatory cytokines IL1β and TNFα. N=4 sham, 3 H/I_Ctrl, 3 H/I_cOE. One-way ANOVA followed by Tukey’s multi-comparison test. IL1β *F*_(2,7)_ = 54.97 *P* < 0.0001, H/I_Ctrl vs. sham *P* = 0.0016, H/I_cOE vs. H/I_Ctrl *P* = 0.0091. TNFα *F*_(2,7)_ = 89.42 *P* < 0.0001, H/I_Ctrl vs. sham *P* = 0.1320, H/I_cOE vs. H/I_Ctrl *P* < 0.0001. **B**, RT-qPCR assays for brain expression of prototypic pro-inflammatory chemokines CCL2 and CXCL10. N=4 sham, 3 H/I_Ctrl, 3 H/I_cOE. One-way ANOVA followed by Tukey’s multi-comparison test. CCL2 *F*_(2,7)_ = 43.80 *P* = 0.0001, H/I_Ctrl vs. sham *P* = 0.0181, H/I_cOE vs. H/I_Ctrl *P* = 0.0029. CXCL10 *F*_(2,7)_ = 23.90 *P* = 0.0007, H/I_Ctrl vs. sham *P* = 0.3901, H/I_cOE vs. H/I_Ctrl *P* = 0.0040. **C-D**, fluorescence immunostaining and quantification of GFAP in the hippocampus at D3 post-H/I injury (Scale bars=100µm). N=5 sham, 4 H/I_Ctrl, 4 H/I_cOE. One-way ANOVA followed by Tukey’s multi-comparison test. *F*_(2,10)_ = 29.03 *P* < 0.0001, H/I_Ctrl vs. sham *P* = 0.0420, H/I_cOE vs. H/I_Ctrl *P* = 0.0030. **E**, RT-qPCR assays for Gfap mRNA levels in the brain at D3 post-H/I injury. N=4 sham, 3 H/I_Ctrl, 3 H/I_cOE. One-way ANOVA followed by Tukey’s multi-comparison test. *F*_(2,7)_ = 22.30 *P* = 0.0009, H/I_Ctrl vs. sham *P* = 0.0272, H/I_cOE vs. H/I_Ctrl *P* = 0.0438. **F**, RT-qPCR quantification for reactive astrocyte marker Vimentin (Vim) in the brain at D3 post-H/I injury. N=4 sham, 3 H/I_Ctrl, 3 H/I_cOE. One-way ANOVA followed by Tukey’s multi-comparison test. *F*_(2,7)_ = 9.509 *P* = 0.0101, H/I_Ctrl vs. sham *P* = 0.5741, H/I_cOE vs. H/I_Ctrl *P* = 0.0452. **G-H**, representative confocal images and quantification showing increased expression of CD68 (microglial activation marker) in the pyramidal layer of hippocampus at D3 post-H/I injury (Scale bars=50µm). N=5 sham, 4 H/I_Ctrl, 4 H/I_cOE. One-way ANOVA followed by Tukey’s multi-comparison test. *F*_(2,10)_ = 341.3 *P* < 0.0001, H/I_Ctrl vs. sham *P* = 0.0089, H/I_cOE vs. H/I_Ctrl *P* < 0.0001. **I**, RT-qPCR quantification of Cd68 in the brain at D3 post-H/I injury. N=4 sham, 3 H/I_Ctrl, 3 H/I_cOE. One-way ANOVA followed by Tukey’s multi-comparison test. *F*_(2,7)_ = 9.630 *P* = 0.0098, H/I_Ctrl vs. sham *P* = 0.7627, H/I_cOE vs. H/I_Ctrl *P* = 0.0320. **J**, RT-qPCR quantification of microglial homeostasis marker Tmem119 in the brain at D3 post-H/I injury. N=4 sham, 3 H/I_Ctrl, 3 H/I_cOE. One-way ANOVA followed by Tukey’s multi-comparison test. *F*_(2,7)_ = 250.3 *P* < 0.0001, H/I_Ctrl vs. sham *P* < 0.0001, H/I_cOE vs. H/I_Ctrl *P* = 0.0415.

Microglia and astroglia play essential roles in brain pro-inflammatory response. To this end, we assessed their activation using molecular and histology assays. Astrocyte activation was significantly enhanced in Serpina3n cOE mice compared with Ctrl mice during H/I injury, as evidenced by elevated levels of GFAP immunoreactive intensity (**Fig. 3C-D**) and mRNA expression (**Fig. 3E**). This activation was further corroborated by the increased expression of Vimentin (**Fig. 3F**), a pan-marker of reactive astrocytes. We also found that SERPINA3N expression significantly increased the expression of CD68, a lysosomal protein upregulated in activated microglia in response to various stimuli, at both histological (**Fig.3G-H**) and molecular levels (**Fig. 3I**) in the hippocampus of Serpina3n cOE mice compared with Serpina3n Ctrl mice after H/I injury. Conversely, SERPINA3N remarkably decreased the expression of TMEM119, a homeostatic microglia marker that is downregulated upon microglial activation (**Fig. 3J**). These results established that SERPINA3N augments the activation of astrocytes and microglia in response to brain injury.

### SERPINA3N promotes neuronal apoptosis and neurodegeneration during neonatal brain injury

HIE causes neurodegeneration and neuronal loss ^31,39^. Consistently, we observed many microinfarct-like lesions after brain H/I injury particularly in the hippocampal regions (**Fig. 4A**) where SERPINA3N was remarkably induced (**Suppl Fig. 9B**). In stark contrast to a neuroprotective role suggested by previous studies, SERPINA3N expression markedly increased the quantity and volume of microinfarct-like lesions in the hippocampus of Serpina3n cOE mice (**Fig. 4A**, arrows). Our quantification showed that the density of NeuN^+^ pyramidal neurons in the hippocampal CA1-CA3 regions was significantly decreased by 50% in Serpina3n cOE mice compared with Serpina3n Ctrl mice (**Fig. 4B**), suggesting that SERPINA3N exerts a deleterious effect on neurodegeneration. In line with the neurodegeneration, we found a significant increase (by 4.5-fold) in the density of cleaved caspase 3 (CC3)-positive cells in hippocampal pyramidal neurons of Serpina3n cOE mice compared with Ctrl mice after H/I injury (**Fig. 4D**). Notably, many CC3^+^ cells exhibited fragmented nuclei with no or scarce NeuN expression (**Fig. 4C and inserts**), suggesting they are degenerating neurons undergoing apoptosis. In addition to neurodegeneration, diffuse axonal injury in the white matter tract is a hallmark pathology in HIE patients. We employed SMI32, a monoclonal antibody recognizing injury-related non-phosphorylated neurofilaments, to identify damaged or dysfunctional axons ^40^. Our results demonstrated that SERPINA3N overexpression aggravated axonal injury as evidenced by significant increase of SMI32 immunoreactive intensity in the subcortical white matter overlaying the hippocampus in Serpina3n cOE mice compared with Serpina3n Ctrl mice (**Fig. 4E-F**). Our findings from genetic overexpression clearly suggest that SERPINA3N promotes neurodegeneration and neuronal loss through apoptosis during brain injury. Our findings weaken previous concept stating that SERPINA3N is neuroprotective in the diseased/injured brain.

**Figure 4.**
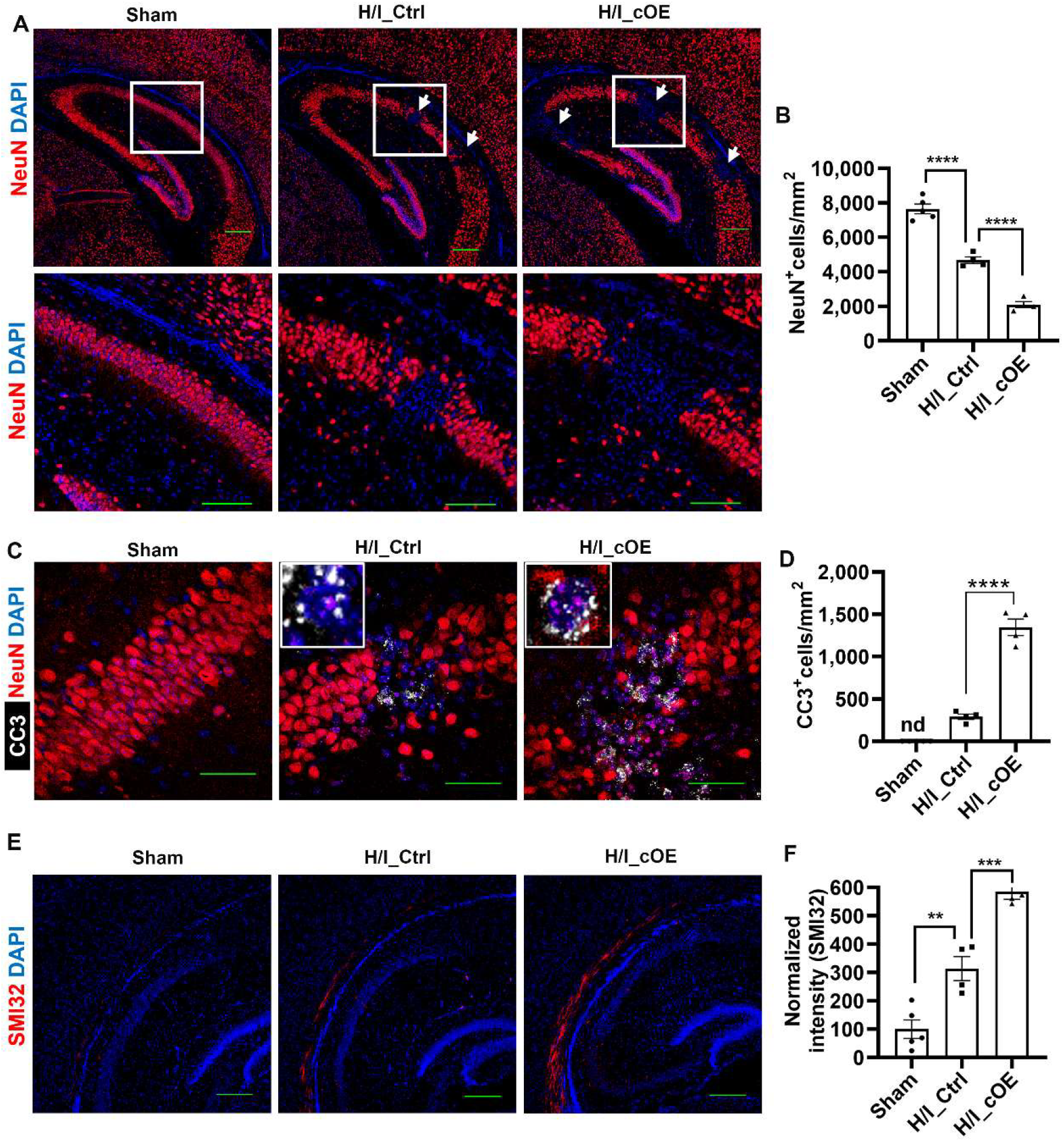
SERPINA3N worsens neuronal death and neurodegeneration during HIE. **A**, fluorescence immunostaining of NeuN depicting the presence of focal lesions (arrows) in the hippocampal CA1-CA3 areas at D3 post-H/I injury. Boxed areas were shown at higher magnification below. **B**, quantification of NeuN^+^ neurons in the pyramidal layer of hippocampus at D3 post-H/I injury. N=5 sham, 4 H/I_Ctrl, 4 H/I_cOE. One-way ANOVA followed by Tukey’s multi-comparison test. *F*_(2,10)_ = 143.0 *P* < 0.0001, H/I_Ctrl vs. sham *P* < 0.0001, H/I_cOE vs. H/I_Ctrl *P* < 0.0001. **C**, immunofluorescence images showing intense signal of cleaved caspase 3 (CC3, canonical apoptosis marker) in the neurodegenerative areas of pyramidal layers at D3 post-H/I injury. Note that CC3^+^ degenerating neurons exhibited fragmented nuclei with sparse NeuN signal (inserts). **D**, quantification of CC3+ cells in the hippocampus at D3 post-H/I injury. N=5 sham, 4 H/I_Ctrl, 4 H/I_cOE. One-way ANOVA followed by Tukey’s multi-comparison test. *F*_(2,10)_ = 168.6 *P* < 0.0001, H/I_cOE vs. H/I_Ctrl *P* < 0.0001. **E**, representative confocal images of low magnification showing increased signal of SMI32, hypo-phosphorylated neurofilaments indicative of axonal injury in the subcortical white matter above the hippocampus. **F**, quantification of SMI32 intensity in the subcortical white matter D3 post-H/I injury. N=5 sham, 4 H/I_Ctrl, 4 H/I_cOE. One-way ANOVA followed by Tukey’s multi-comparison test. *F*_(2,10)_ = 50.98 *P* < 0.0001, H/I_Ctrl vs. sham *P* = 0.0032, H/I_cOE vs. H/I_Ctrl *P* = 0.0008. Scale bars: A, 200µm (low magnification), 100µm (high magnification), C, 50µm, E, 200µm.

### SERPINA3N inhibits oligodendrocyte differentiation and disturbs developmental myelination during neonatal brain injury

Oligodendrocyte differentiation and developmental myelination are disrupted in HIE patients and preclinical animal models ^32,41^. We quantified the population of OPCs and differentiated OLs in the subcortical white matter where diffuse axonal injury was observed. The density of SOX10^+^PDGFRa^+^ OPCs (**Fig. 5A**) in Serpina3n cOE mice was similar to that in Serpina3n Ctrl mice at day 3 post-H/I injury (**Fig. 5B**). Yet, SERPINA3N expression significantly diminished the population of SOX10^+^CC1^+^ OLs (**Fig. 5C**) in Serpina3n cOE mice compared with Ctrl mice at day 3 post-H/I injury (**Fig. 5D**). These data suggest that SERPINA3N worsens the disturbance of OPC differentiation into OLs under brain injury conditions. Consistent with the diminished oligodendrocyte population, developmental myelination, visualized by MBP immunoreactivity (**Fig. 5E**), remarkably decreased in the subcortical white matter of H/I-injured Serpina3n cOE mice compared with H/I-injured Ctrl mice (**Fig. 5F**). Taken together, our results indicate that SERPINA3N inhibits oligodendrocyte differentiation and disrupts developmental myelination under injury conditions though it is dispensable for normal oligodendroglial development under homeostasis.

**Figure 5.**
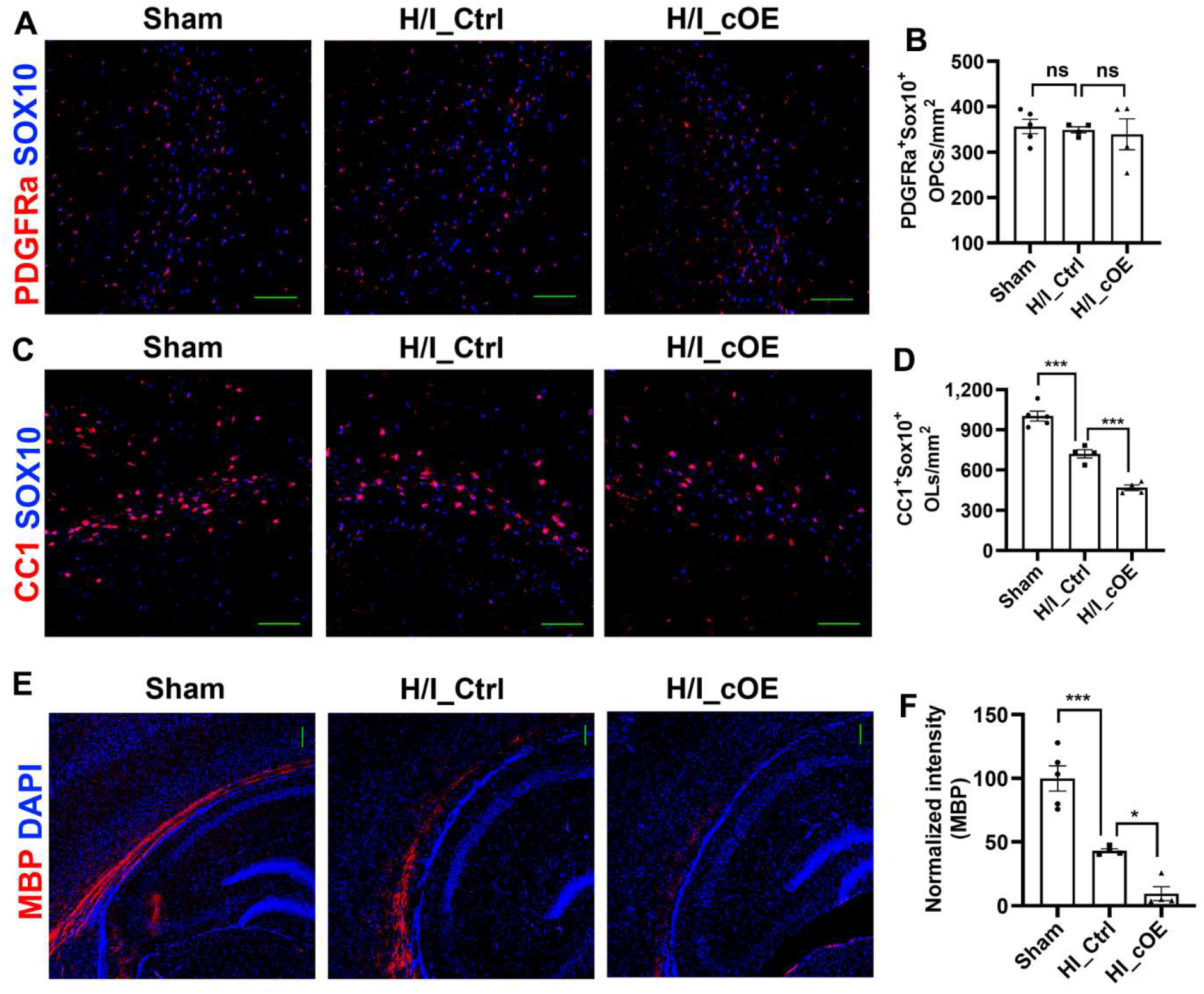
SERPINA3N inhibits myelination and oligodendroglial differentiation during HIE. **A-B**, fluorescence immunostaining and quantification of SOX10 and PDGFRa (OPC marker) in the subcortical white matter overlaying the hippocampus at D3 post-H/I injury. N=5 sham, 4 H/I_Ctrl, 4 H/I_cOE. One-way ANOVA followed by Tukey’s multi-comparison test. *F*_(2,10)_ = 0.1682 *P =* 0.8520, H/I_Ctrl vs. sham *P* = 0.9689, H/I_cOE vs. H/I_Ctrl *P* = 0.9476. **C-D**, representative confocal images and quantification of CC1^+^SOX10^+^ mature OLs in the subcortical white matter overlaying the hippocampus at D3 post-H/I injury. N=5 sham, 4 H/I_Ctrl, 4 H/I_cOE. One-way ANOVA followed by Tukey’s multi-comparison test. *F*_(2,10)_ = 74.09 *P* < 0.0001, H/I_Ctrl vs. sham *P* = 0.0002, H/I_cOE vs. H/I_Ctrl *P* = 0.0008. **E-F**, fluorescence immunostaining of myelin basic protein (MBP, a myelin marker) in the subcortical white matter overlaying the hippocampus at D3 post-H/I injury. N=5 sham, 4 H/I_Ctrl, 4 H/I_cOE. One-way ANOVA followed by Tukey’s multi-comparison test. *F*_(2,10)_ = 41.43 *P* < 0.0001, H/I_Ctrl vs. sham *P* = 0.0006, H/I_cOE vs. H/I_Ctrl *P* = 0.0266. Scale bars: A, 100µm, C, 100µm, E, 100µm

## Discussion

SERPINA3N/SERPINA3 has been gaining increasing attention since the discovery of its deposit in β-amyloid plaques in Alzheimer’s disease in 1988 ^7^. Recently, this molecule has been identified as a top signature gene upregulated in disease-associated neural cells of various neurological diseases and injuries ^42-45^. However, its role in regulating CNS development and pathophysiology remains elusive^2^. Using non-invasive genetic approaches to express SERPINA3N specifically in brain neural cells, this study made several significant findings: 1) enforced gain-of-function in neural precursor cells, which express otherwise undetectable levels of SERPINA3N, does not affect neurodevelopment or improve animal cognition; 2) SERPINA3N does not cause BBB abnormalities in the homeostatic brain yet it promotes BBB disruption after brain injury; 3) SERPINA3N drives pro-inflammatory responses to brain injury and enhances glial activation; 4) SERPINA3N aggravates neurodegeneration through apoptosis-mediated neuronal death and disrupts normal oligodendrocyte development and myelination under pathological conditions. Our findings suggest that future interventions should inhibit, rather than augment, SERPINA3N to achieve therapeutic effects on brain diseases and injuries.

Enforced expression of human SERPINA3 was reported to enhance neurogenesis and improved cognitive ability in mice ^3^. SERPINA3 is highly expressed in human NPCs whereas the murine homolog SERPINA3N is downregulated to undetectable levels in mice ^3^. From an evolutionary perspective, it is attempting to hypothesize that SERPINA3/SERPINA3N is a candidate molecule that drives neocortical folding in human brain. If this hypothesis holds true, one can expect that re-gaining SERPINA3N in Serpina3n-negative murine NPCs will similarly promote neurogenesis and cognitive ability in mice. In line with the report ^3^, we found that SERPINA3N gain-of-function does not affect gliogenesis, nor does it lead to abnormal neurobehaviors in mice. However, our findings - SERPINA3N gain-of-function does not improve cognitive function - are quite surprising. The unaltered neurobehavior is consistent with the normal development of neurons and glial cells in Serpina3n cOE mice. Our results do not necessarily dispute previous conclusion ^3^. These discrepant findings may suggest a possible functional difference between human SERPINA3N and murine SERPINA3N in regulating neurogenesis and mouse cognition given that they share ∼61% homology in the primary peptide sequence ^2^ despite their functional similarity ^1^. Nevertheless, our study established that murine SERPINA3N is dispensable for homeostatic brain development and neurobehavior ^6^. Future studies are needed to identify the counterpart of human SERPINA3 among murine Serpina3 cluster members ^1^ in promoting neurogenesis and cognitive function in mice.

A recent study reported that exogenously administered SERPINA3N is sufficient to cause vascular inflammation and BBB disruption in the homeostatic mouse brain ^22^. Our results do not support this conclusion. We found that brain overexpression of SERPINA3N by the non-invasive *Cre-loxP* genetic approach does not cause BBB disruption, neuroinflammation, or glial activation during brain homeostatic development. We reasoned that the phenotypes observed in that study ^22^ are likely due to traumatic brain injury created by invasive stereotaxic injections. Indeed, we have shown that SERPINA3N promotes BBB disruption and vascular changes in response to brain injury. The data from previous studies ^6,22^ and the current study altogether suggest that SERPINA3N is an injury-induced factor that plays crucial roles only under diseased/injured conditions. Importantly, our data highlighted the superiority of using non-invasive *Cre-loxP* genetic approaches to invasive approaches (such as exogenous injections) to study the biological role of SERPINA3N in regulating certain biological processes.

BBB dysfunction, neuroinflammation, and neurodegeneration are shared pathophysiology in various CNS neurological diseases/injuries, such as hypoxic-ischemic brain injury, cerebral stroke, CNS trauma, and neurodegenerative disorders. Recent studies using invasive approaches concluded that viral vector injection-mediated SERPINA3N expression attenuated BBB damage, neuroinflammation, and neuronal apoptosis in response to brain ischemic damage ^16,17^ and trauma ^19^, pointing to a protective effect on cerebral ischemic pathophysiology. However, the protective role is difficult to reconcile with the clinical observations that higher SERPINA3 level was significantly associated with worsened vascular and neural symptoms in patients of cerebral ischemic stroke^13^ and intracerebral hemorrhage ^9^. In line with the clinical studies, our results show that SERPINA3N aggravates brain injury, as evidenced by worsened brain pathology of BBB disruption, neuroinflammation, and neurodegeneration in response to hypoxic-ischemic brain injury in Serpina3n cOE mice. Our conclusion was made by leveraging the non-invasive *Cre-loxP* approaches to express SERPINA3N specifically in neural cells whereas others ^16,17,19^ employed viral vector-mediated SERPINA3N overexpression. We cannot exclude the possibility that SERPINA3N regulation of the shared brain pathophysiology is different in HIE from that in ischemic stroke or brain trauma. Alternatively, SERPINA3N expression by peripheral organs and/or brain non-neural cells may play a crucial role in regulating brain pathology under diseased/injured conditions. In this regard, it is important to determine whether brain pathology is aggravated or diminished using our neural cell-specific Serpina3n cOE mouse models after cerebral ischemia or brain trauma.

The role of SERPINA3N in oligodendrocyte differentiation and myelination after brain injury remains elusive. Our findings demonstrate for the first time that SERPINA3N upregulation disturbs oligodendrocyte lineage maturation and leads to hypomyelination after hypoxia-ischemia brain injury, suggesting that inhibiting SERPINA3N is of therapeutic effect on insufficient myelination in HIE survivors. Our results show that SERPINA3N expression does not affect OPC population, instead, it inhibits the differentiation of OPCs into mature OLs. The mechanism underlying SERPINA3N-regulated hypomyelination remains elusive and it may involve in SERPINA3N-mediated OL viability during brain injury as suggested by our previous *in vitro* data ^6^, which ultimately leads to fewer OLs for subsequent myelination. Assessing OL cell death and viability in injured Serpina3n cOE mice may provide new insight into this possible mechanism.

Taken together, by leveraging the powerful, non-invasive expression of SERPINA3N in brain neural cells, the current study challenges the prevailing paradigm of “SERPINA3N neuroprotection” after brain diseases and injuries. Our results support the neurotoxic role of SERPINA3N in BBB dysfunction, neuroinflammation, neuronal loss and axonal injury, and hypomyelination. The findings provide instructions for future designs of dampening SERPINA3N expression and/or function to achieve therapeutic benefits.

### Limitations of the study

First, we used *Cre-loxP* tools to drive SERPINA3N expression in NPCs of embryos and NPC-generated neural progenies of postnatal mice. This study is yet to answer the question whether individual neural cell type-derived SERPINA3N exerts similar or distinct effects on shared brain pathophysiology. Second, we tested the role of SERPINA3N in modulating the shared brain pathology by using neonatal brain hypoxic-ischemic brain injury. Assessing injury type-specific role of SERPINA3N using our Serpina3n cOE models is important to draw a general conclusion interpreting SERPINA3N’s function in the diseased CNS. Third, SERPINA3N is elevated after brain injury, it would be equally informative to employ brain-specific *Serpina3n* loss-of-function to strengthen our conclusions.

## Materials and Methods

### Transgenic mice

All transgenic mice were maintained on a C57BL/6 background under a 12 h light/dark cycle with ad libitum access to food and water. Both male and female mice were included in this study. Four transgenic lines were used. *Nestin-Cre* (RRID:IMSR_JAX:003771) and *Serpina3n-floxed* (RRID:IMSR_JAX:027511) mice obtained from The Jackson Laboratory. The *Rosa26*^*CAG-Serpina3n-P2A-EGFP*^ knock-in line and the *Serpina3n-tdTom* reporter line were generated by the Guo laboratory.

*The CAG-Serpina3n-P2A-EGFP* construct was knocked in the mouse endogenous Rosa26 locus (**Suppl Fig. 1**) using CRISPR/Cas9-mediated Extreme Genome Editing (EGE). A single sgRNA was designed to target the intron 1–2 region of the mouse *Rosa26* locus. The donor construct was assembled in the TV-4G backbone and consisted of a ∼1.5 kb 5′ homology arm, a CAG promoter, a floxed STOP cassette (*LoxP-STOP-LoxP*), the mouse Serpina3n coding sequence fused to a *P2A-EGFP* tag, followed by a WPRE-polyadenylation termination sequence, and a ∼1.5 kb 3′ homology arm. Cas9 mRNA, the sgRNA, and the targeting vector were co-microinjected into C57BL/6J-derived zygotes, which were subsequently implanted into pseudopregnant recipient females. The “*CAG promoter–LoxP–STOP–LoxP–Serpina3n–P2A– EGFP–WPRE–pA*” cassette was successfully integrated into the intron 1-2 region of the Rosa26 safe-harbor locus. Out of 572 zygotes transferred, 100 live pups were obtained, of which 6 were identified as F0 founders carrying the correct knock-in allele. These F0 founders were crossed with wild-type C57BL/6J mice to generate F1 offspring. Germline transmission was confirmed in heterozygous F1 mice by PCR, Sanger sequencing, and Southern blot analysis using probes at both the 5′ and 3′ ends to exclude random integration events. PCR amplification of genomic DNA was performed using the following primer sets: LSL-Serpina3n-GT-F: AGTCGCTCTGAGTTGTTATCAG; LSL-Serpina3n-GT-R: TGAGCATGTCTTTAATCTACCTCGATG; LSL-Serpina3n-PCR-R: AGTCCCTATTGGCGTTACTATGG (see Suppl Fig. 1).

A Serpina3n-tdTomato knock-in (KI) mouse line was generated using CRISPR/Cas9-based EGE technology (Suppl Fig. 9). A single sgRNA was designed to target exon 5 of the mouse *Serpina3n* gene. The donor construct was assembled in the TV-4G vector and contained a ∼1.5 kb 5′ homology arm, exons 4-5 fused to a *P2A-tdTomato* cassette, and a ∼1.5 kb 3′ homology arm. Cas9 mRNA, the sgRNA, and the targeting vector DNA were microinjected into zygotes derived from C57BL/6J mice, and injected embryos were transferred into pseudopregnant surrogate females. The *P2A-tdTomato* cassette was precisely inserted into exon 5, immediately upstream of the 3′ untranslated region (3′UTR) of Serpina3n. A total of 537 zygotes were transferred, resulting in 88 live pups, of which 7 F0 mice were identified as positive founders. These F0 founders were bred with wild-type C57BL/6J mice to generate F1 offspring. Germline transmission in heterozygous F1 animals was confirmed by PCR genotyping, DNA sequencing, and Southern blot analysis using probes at both the 5′ and 3′ ends to exclude random integration events. PCR amplification of genomic DNA was performed using the following three primer sets: EGE-YQL-0142-A-WT-F: TCCCTGGTCTCCACTGAGTGATGTG; EGE-YQL-0142-A-WT-R: GCTACCAAGAAGTCTGGGAGCATGG; and tdtomato-15R:

AACTCTTTGATGACCTCCTCG. The primer pair of EGE-YQL-0142-A-WT-F/ EGE-YQL-0142-A-WT-R produced a PCR amplicon of 435 bp in wildtype mice whereas the primer pair of EGE-YQL-0142-A-WT-F/tdTomato-15R generated a PCR amplicon of 333 bp in mutant mice.

Animal genotype was determined by PCR of genomic DNA extracted from tail tissue. All Cre lines were maintained as heterozygosity. All animals and procedures were approved by UC Davis IACUC.

### Hypoxia/ischemia injury

Neonatal H/I injury was induced in P6 mice pups according to previously published protocols ^33^. Briefly, pups were anesthetized by indirect cooling on wet ice, followed by permanent occlusion of the right common carotid artery through electrocauterization. The pups were kept on a thermal blanket until fully awake, then returned with the dam. After one hour of recovery, hypoxia was induced by placing the pups in a sealed chamber infused with nitrogen to maintain 6.0% O_2_ for 45 min. Following hypoxia, pups were housed with the dam until further experiments.

### Behavioral analysis

#### Barnes Maze Assessment

Spatial learning, memory, and locomotor activity were assessed using the Barnes maze. The apparatus consisted of a circular platform (100 cm in diameter) containing 20 evenly spaced holes (10 cm in diameter), one of which connected to an escape box. Mice were placed in the center of the platform and required to locate the goal box. Experiments were conducted in a quiet room with distinct visual cues positioned around the maze. Animals underwent training for 5 consecutive days, followed by probe testing on day 6. Each trial lasted a maximum of 5 minutes, unless the animal entered the goal box earlier. Behavioral parameters, specifically latency to goal entry, were recorded and analyzed using EthoVision XT 14 software.

#### Open field test

Animals were acclimated to the behavioral testing room for 30 minutes before testing. Mouse behavior was then recorded for 10 minutes in an arena measuring 42 × 42 × 37 cm. Parameters analyzed included time spent in the center zone (25% of the arena), time spent in the border zone, total distance traveled, and average speed. Data was recorded and analyzed using EthoVision XT 14 software.

#### Elevated plus maze

Exploratory behavior was assessed for 5 minutes in the elevated plus maze, consisting of two open arms and two closed arms elevated 50 cm above the floor. Each arm measured 50 × 10 cm, with closed arms enclosed by 38 cm high walls, and a central area of 10 × 10 cm. An overhead camera provided a full view of the maze and was connected to a computer running EthoVision XT 14 for automated tracking. Parameters analyzed included time spent in the open and closed arms, frequency of entries into open and closed arms, average speed, and total distance travel.

#### Tissue processing and immunohistochemistry

Mice were deeply anesthetized with an IP injection of a ketamine/xylazine mixture and transcardially perfused with ice-cold PBS, followed by freshly prepared 4% PFA. Brains were harvested and post-fixed in 4%PFA for 5 hr at RT, cryoprotected in 30% sucrose for 24 hr at 4°C, embedded in OCT medium (Fisher Healthcare), and kept frozen at −80°C until 14 μm of serial coronal sections were cut using a cryostat (model CM1900-3-1, Leica). Sections were blocked for 1.5 hr in the blocking solution containing 10% donkey serum in PBST (PBS with 0.1% Tween-20), then incubated overnight at 4°C with the following primary antibodies: Rabbit anti-NeuN (1:400, #A0951, Abclonal); Mouse anti-NeuN (1:400, #MAB377, Millipore); Rabbit anti-Iba1 (1:200, #019-19741, WAKO); Mouse anti-GFAP (1:20000, #MAB360, Millipore); Mouse anti-MBP (1:200, #NE1019, Millipore Sigma); Rabbit anti-Ki67(1:100, #9129, Cell Signalling Technology); Rabbit anti-RFP(1:100, #600-401-379, Rockland); Rabbit anti-TCF7L2 (1:200, #2569, Cell Signalling Technology): Rat anti-CD68 (1:200, #MCA1957, Bio-Rad), Mouse anti-CC1 (1:100, #OP80, Millipore Sigma), Rabbit anti-SOX10 (1:200, #ab155279, Abcam), Goat anti-PDGFRα (1:200, #AF1062, NOVUS Biologicals), Rabbit anti-Laminin (1:150, NB300-144, NOVUS), Goat anti-Albumin (1:400, #A90-134A, FORTIS), Mouse anti-Claudin 5 (1:200, #35-2500, Thermo Fisher), mouse antiSMI32 (1:200, #801706, Biolegend), Rabbit anti-cleaved Caspase 3 (1:100, #9661, Cell Signalling Technology). After washing three times in PBST, the sections were incubated with the appropriate Alexa Fluor-conjugated secondary antibodies (Jackson Immuno Research Laboratories) for 1.5 hr at RT. An additional three washes were performed, and the air-dried sections were mounted with antifading mounting medium after DAPI nuclear staining.

#### Total RNA extraction, cDNA preparation, and RT-qPCR

The brain was dissected and lysed with QIAzol (QIAGEN). Total RNA was extracted using the RNeasy Plus Kit (Cat#74034, QIAGEN) following the manufacturer’s instruction. RNA quality and concentration were measured by NanoDrop 2000 spectrophotometer (Thermo Fisher Scientific), and cDNA was reverse transcribed using ABScript II kit (Cat #RK20400, ABclonal). Quantitative PCR assays were performed using gene-specific primers and SYBR green-based master mix (Cat# RK21203, ABclonal) on a real-time PCR system (AriaMx, Agilent). The relative gene expression level was calculated using the equation 2^ (Ct of HSP90-Ct of target gene), where HSP90 serves as a reference gene. Primers used for the RT-qPCR:

Sox10: ACACCTTGGGACACGGTTTTC/TAGGTCTTGTTCCTCGGCCAT, Plp1: GTTCCAGAGGCCAACATCAAG/CTTGTCGGGATGTCCTAGCC, Mbp: GGCGGTGACAGACTCCAAG/GAAGCTCGTCGGACTCTGAG, Mobp: AACTCCAAGCGTGAGATCGT/CAGAGGCTGTCCATTCACAA, Myrf: CAGACCCAGGTGCTACAC/TCCTGCTTGATCATTCCGTTC, Iba1: GGACAGACTGCCAGCCTAAG/GACGGCAGATCCTCATCATT, Mog: AGCTGCTTCCTCTCCCTTCTC/ACTAAAGCCCGGATGGGATAC, Gfap: GTGTCAGAAGGCCACCTCAAG/CGAGTCCTTAATGACCTCACCAT, Aldh1l1: AGCCACCTATGAGGGCATTC/TGAGTGTCGAGTTGAAAAACGTC, Tmem119: CCTACTCTGTGTCACTCCCG/CACGTACTGCCGGAAGAAATC, Cx3cr1: GAGTATGACGATTCTGCTGAGG/CAGACCGAACGTGAAGACGAG, Cd68: CCATCCTTCACGATGACACCT/GGCAGGGTTATGAGTGACAGTT, NeuN: ATCGTAGAGGGACGGAAAATTGA/GTTCCCAGGCTTCTTATTGGTC, Tubb3: TAGACCCCAGCGGCAACTAT/GTTCCAGGTTCCAAGTCCACC, Syp: CAGTTCCGGGTGGTCAAGG/ACTCTCCGTCTTGTTGGCAC, Glo1: CCCTCGTGGATTTGGTCACA/AGCCGTCAGGGTCTTGAATG, IL1β: GCACTACAGGCTCCGAGATGAA/GTCGTTGCTTGGTTCTCCTTGT, Tnfα: GACGTGGAACTGGCAGAAGAG/TTGGTGGTTTGTGAGTGTGAG, Ccl2: CAGCAAGATGATCCCAATGA/TCTGGACCCATTCCTTCTTG, Cxcl10: GCTGCAACTGCATCCATATC/GGATTCAGACATCTCTGCTCAT, Vim: GAGGAGATGAGGGAGTTGCG/CTGCAATTTTTCTCGCAGCC, HSP90: AAACAAGGAGATTTTCCTCCGC/CCGTCAGGCTCTCATATCGAAT

#### Protein preparation and Western blot

Brain tissues were homogenized in N-PER Neuronal Protein Extraction Reagent (Thermo Fisher Scientific) supplemented with protease and phosphatase inhibitor cocktails. The homogenates were kept on ice for 30 minutes and then clarified by centrifugation at 14,000 × rpm for 30 minutes at 4 °C. Protein concentration was quantified using the BCA assay kit (Thermo Fisher Scientific). Equal protein amounts (30 µg per sample) were separated by SDS–PAGE on AnykD Mini-PROTEAN TGX precast gels (Bio-Rad) and subsequently transferred onto 0.2 µm nitrocellulose membranes (Bio-Rad) using the Trans-Blot Turbo Transfer System (Bio-Rad). Membranes were blocked for 5 min at room temperature with EveryBlot Blocking Buffer (Bio-Rad Cat. #12010020) and then incubated with primary antibodies overnight at 4 °C. After washing, membranes were treated with HRP-conjugated secondary antibodies, and signals were detected using Western Lightning Plus ECL (PerkinElmer). Band densities were quantified with NIH ImageJ software. Antibodies used for Western blot are listed below: Goat polyclonal anti-Serpina3n (1:1000, R&D System, Cat# AF4709; RRID:AB_2270116), Rabbit polyclonal anti-IBA1 (1:1000, WAKO, Cat# 019–19741; RRID:AB_839504), Rabbit polyclonal anti-GFAP (1:1000, Millipore, Cat# MAB36014; RRID:AB_11212597), Mouse Monoclonal anti-MOG (1:1000, Invitrogen, Cat# MA5-24645; RRID:AB_2637260), Mouse Monoclonal anti-Claudin 5 (1:1000, Invitrogen, Cat# 35-2500, AB_2533200), Rabbit Monoclonal anti-VCAM-1(1:1000, Invitrogen, Cat# MA5-31965; RRID:AB_2809259), Rabbit monoclonal anti-GAPDH (1:1000, Cell Signaling Technology Cat# 2118, RRID:AB_561053). Secondary antibodies conjugated to HRP were sourced from Thermo Fisher Scientific.

#### EdU labelling

P0 and P14 mice were injected with 50mg/kg EdU and their brains were harvested 2 hr and 24 hr after injection, respectively. For the embryonic study, timed pregnant mouse at E15.5 were intraperitoneally injected with 50mg/kg of EdU 2hr before tissue harvesting. The harvested brains were fixed in 4%PFA for 5 hr, cryoprotected in 30% sucrose for 24 hr, and cut into 14 µm sagittal sections using a cryostat (model CM1900-3-1, Leica). Incorporation of EdU into DNA was detected using Click-iT Plus EdU Cell Proliferation Kit with Alexa Fluo 647 (Cat. #C10640, Thermo Fisher). The sectioned were co-stained with rabbit anti-NeuN antibody (1:400, #A0951, Abclonal) and DAPI, then imaged using a Nikon A1 confocal microscope.

#### Data acquisition and statistical analysis

All experiments included both male and female mice. Experimental group identity (genotype and treatment) was blinded to investigators performing data acquisition and quantification. Results are displayed as mean ± standard error of mean (s.e.m.). Scatter dot plots are presented throughout the study, with each point corresponding to a single animal or an independent replicate. For two-group comparisons, unpaired two-tailed Student’s t-test was used. For analyzing three groups, one-way ANOVA was employed, followed by Tukey’s post hoc test for multiple comparisons. All statistical testing and visualization were carried out in GraphPad Prism (version 8.0). Statistical significance was defined as p-value < 0.05. F values (ANOVA), t values (Student’s t test), degree of freedom, and exact P-values were presented in the figure legends. Significance levels are represented in figures as follows: ∗p < 0.05, ∗∗p < 0.01, ∗∗∗p < 0.001; “ns” indicates not significant (p > 0.05).

## Acknowledgements

We thank the funding agencies of NIH (R21NS125464, R01NS123080, R01NS123165, R01NS134887 to FG) and Shriners Hospitals for Children (85101-NCA-22, 85113-NCA-23 to FG, 84312-NCA-24 to MZ) for supporting the work.

