## Supplemental Figure 1 to Figure 9 for "A paradigm shift of SERPINA3N in neurobehavioral development and brain injury"

### Supplemental Data

### Nine Supplemental Figures and Legends

### Suppl Fig 1 – generation of loxP-STOP-loxP-Serpina3n transgenic mice

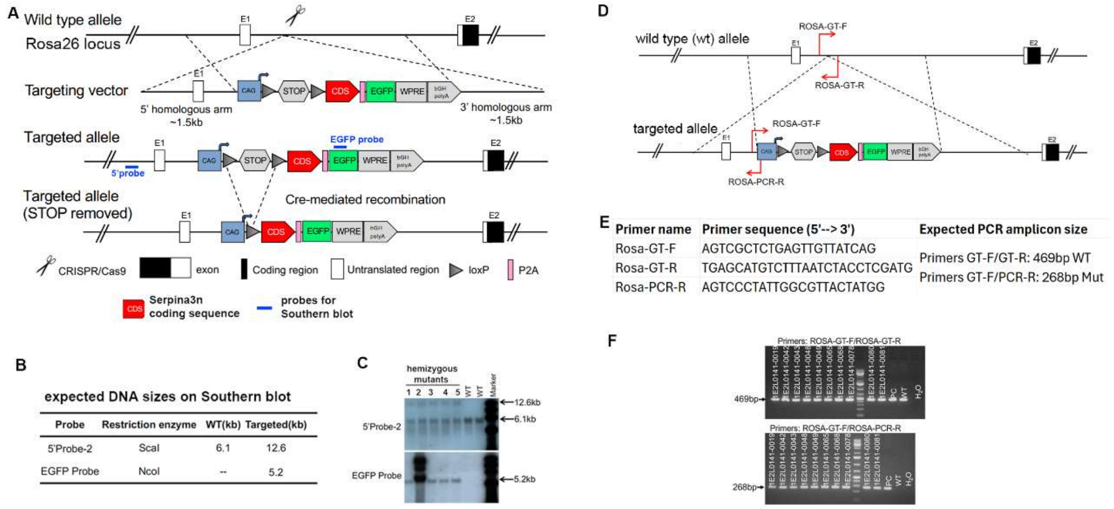


**A**, schematic diagram showing the targeting vector, targeted allele, and mutant allele after Cre-mediated deletion of the STOP codon.

**B-C**, Southern blot assay of mouse genomic DNA showing expected band sizes in the targeted allele.

**D-E**, genotyping PCR primer locations and sequences and expected PCR amplicon sizes.

**F**, representative genotyping results of wild type and mutant alleles.

**Suppl Fig 2 – neuronal and glial cell development is normal at weaning ages in SERPINA3N-expressing transgenic mice**

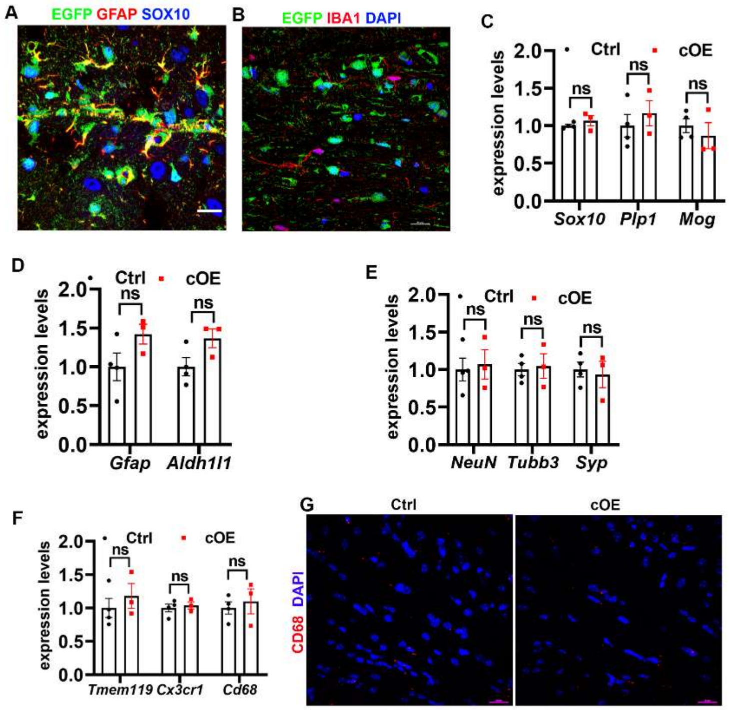


**A**, fluorescence immunostaining showing that EGFP is expressed in GFAP$^+$ astroglial lineage cells and SOX10$^+$ oligodendroglial lineage cells in the brain of Serpina3n cOE mice at weaning ages. Scale bar=10μm.

**B**, fluorescence immunostaining showing that EGFP is absent from IBA1$^+$ microglia (myeloid lineage) in the brain of Serpina3n cOE mice at weaning ages. Scale bar=10μm.

**C**, RT-qPCR assays of oligodendrocyte markers Sox10, Plp1, and Mag in the brain of P22 mice. N=4 Ctrl, 3 cOE, unpaired Student's t test, $t_{(5)}$ = 1.783 *P* = 0.1347 Gfap, $t_{(5)}$ = 2.129 *P* = 0.0865 Aldh1l1.

**D**, RT-qPCR assays of astrocyte markers Gfap and Aldh1l1 in the brain of P22 mice. N=4 Ctrl, 3 cOE, unpaired Student's t test, $t_{(5)}$ = 1.090 *P* = 0.3255 Sox10, $t_{(5)}$ = 0.7310 *P* = 0.4976 Plp1, $t_{(5)}$ = 0.7499 *P* = 0.490 Mag.

**E**, RT-qPCR assays of neuronal markers NeuN, Tubb3, and Synapsin (Syp) in the brain of P22 mice. N=4 Ctrl, 3 cOE, unpaired Student's t test, $t_{(5)}$ = 0.2858 *P* = 0.7865 NeuN, $t_{(5)}$ = 0.2797 *P* = 0.7909 Tubb3, $t_{(5)}$ = 0.3522 *P* = 0.7391 Syp.

**F**, RT-qPCR assays of microglial homeostasis markers Tmem119 and Cx3cr1 and activation marker Cd68 in the brain of P22 mice. N=4 Ctrl, 3 cOE, unpaired Student's t test, $t_{(5)}$ = 0.7871 *P* = 0.4669 Tmem119, $t_{(5)}$ = 0.4660 *P* = 0.6608 Cx3cr1, $t_{(5)}$ = 0.5195 *P* = 0.6256 Cd68.

**G,** fluorescence immunostaining showing no microglial activation in the brain of Serpina3n cOE mice at weaning ages. Scale bar=10μm.

**Suppl Fig 3 - Serpina3n is dispensable for neurobehavioral development in the adult animals**

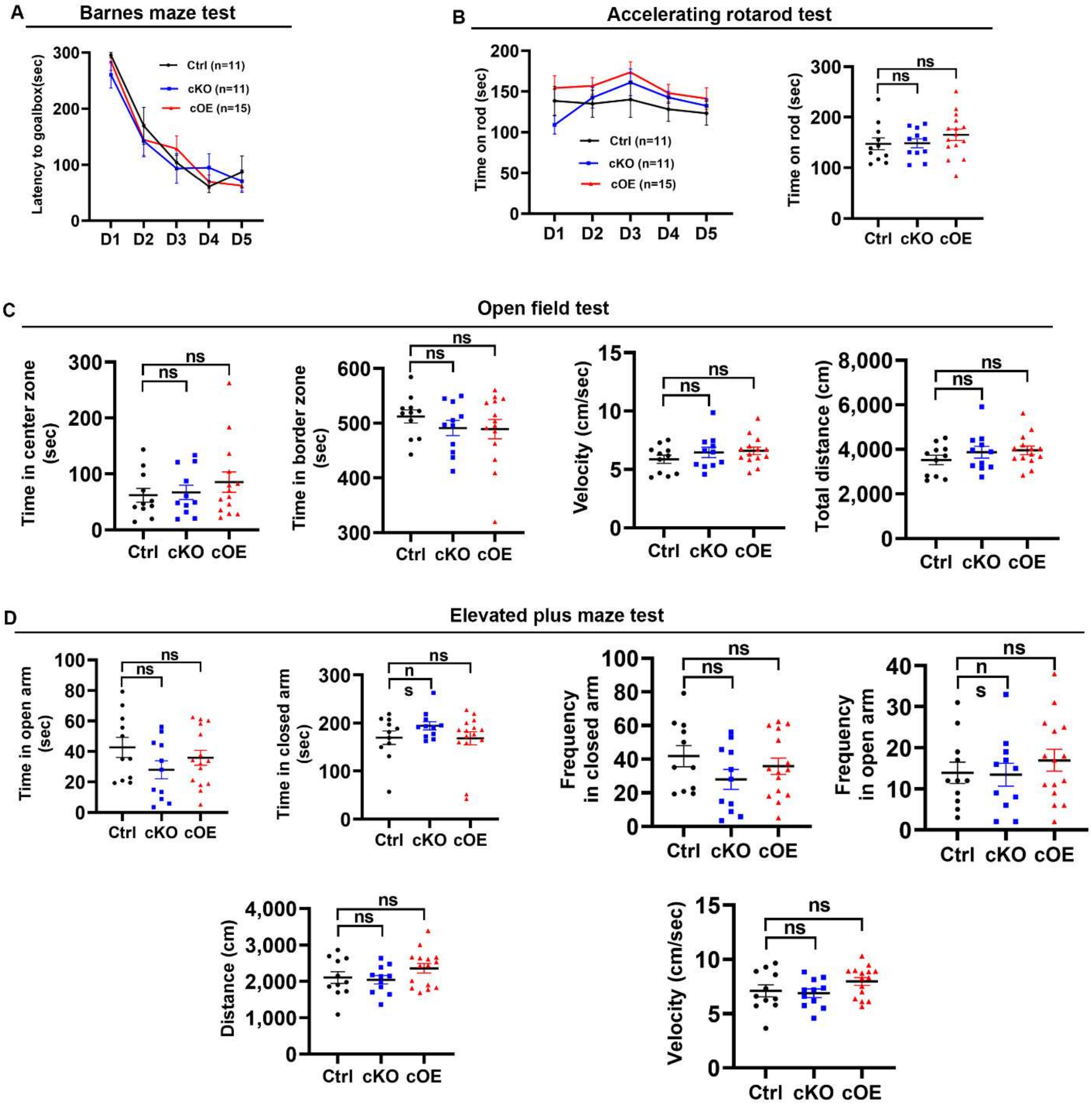


**A,** Time latency (seconds) to find goalbox during the training sessions (time course from day 1 D1 to D5) of Barnes maze test. Each session takes 5 minutes. Two-month-old mice were used for testing. N=11 Ctrl, 11 cKO, 15 cOE. Two-way ANOVA repeated measures, time course $F_{(4, 136)}$ = 57.19 $P$ < 0.0001, genotype $F_{(2, 34)}$ = 0.1786 $P$ = 0.8372.

**B**, Time on rod (seconds) of accelerating rotarod test during the training session (D1-D5, left) and test session (D6). N=11 Ctrl, 11 cKO, 15 cOE. One-way ANOVA, $F_{(2, 34)}$ = 0.9291 $P$ = 0.4075.

**C**, Time in center or border zones (indicating anxiety levels) and moving velocity and distance (indicating locomotion activity) in the open field test (10 minutes). N=11 Ctrl, 11 cKO, 14 cOE. One-way ANOVA, $F_{(2, 33)}$ = 0.6681 $P$ = 0.5195 time in center, $F_{(2, 33)}$ = 0.6724 $P$ = 0.5173 time in border, $F_{(2, 33)}$ = 1.064 $P$ = 0.3566 velocity, $F_{(2, 33)}$ = 1.062 $P$ = 0.3574 distance.

**D**, Time and frequency in open and closed arms (indicating anxiety levels) and moving velocity and distance (indicating locomotion activity) in the elevated plus maze test (5 minutes). N=11 Ctrl, 11 cKO, 15 cOE. One-way ANOVA, $F_{(2, 34)}$ = 1.485 $P$ = 0.2408 time in open arm, $F_{(2, 34)}$ = 1.233 $P$ = 0.3404 time in closed arm, $F_{(2, 34)}$ = 0.5277 $P$ = 0.5947 frequency in open arm, $F_{(2, 34)}$ = 1.370 $P$ = 0.2679 frequency in closed arm, $F_{(2, 34)}$ = 1.610 $P$ = 0.2147 distance traveled, $F_{(2, 34)}$ = 1.887 $P$ = 0.1670 moving velocity.

**Suppl Fig 4 - indistinguishable levels of glial cell marker gene expression in the adult brain**

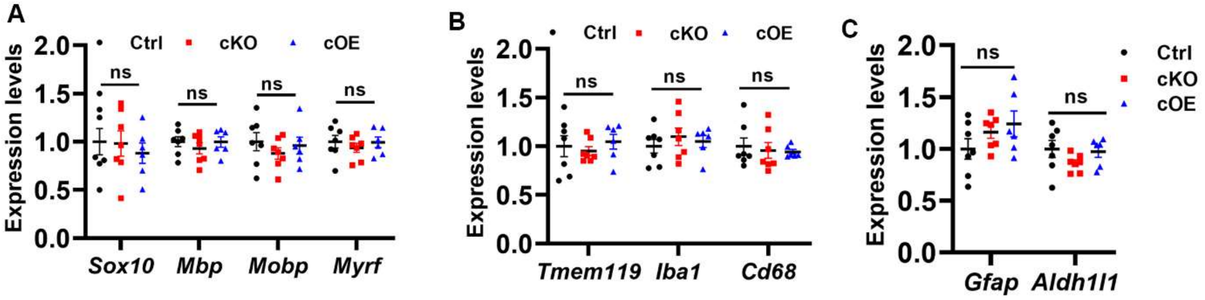


**A**, RT-qPCR assays for oligodendrocyte markers Sox10, Mbp, Mobp, and Myrf in the brain of adult mice. N=7 Ctrl, 7 cKO, 6 cOE. One-way ANOVA, $F_{(2,17)}$ = 0.2390 $P$ = 0.7900 Sox10, $F_{(2,17)}$ = 0.5353 $P$ = 0.5950 Mbp, $F_{(2,17)}$ = 0.6144 $P$ = 0.5525 Mobp, $F_{(2,17)}$ = 0.3993 $P$ = 0.6769 Myrf.

**B**, RT-qPCR assays for microglial markers Tmem119, Iba1, and Cd68 in the brain of adult mice. N=7 Ctrl, 7 cKO, 6 cOE. One-way ANOVA, $F_{(2,17)}$ = 0.3404 $P$ = 0.7163 Tmem119, $F_{(2,17)}$ = 0.4296 $P$ = 0.6576 Iba1, $F_{(2,17)}$ = 0.1742 $P$ = 0.8416 Cd68.

**C**, RT-qPCR assays for astrocyte markers Gfap and Aldh1l1 in the brain of adult mice. N=7 Ctrl, 7 cKO, 6 cOE. One-way ANOVA, $F_{(2,17)}$ = 1.673 $P$ = 0.2171 Gfap, $F_{(2,17)}$ = 1.515 $P$ = 0.2481 Aldh1l1.

**Suppl Fig 5 - Serpina3n expression does not affect neural precursor cell proliferation**

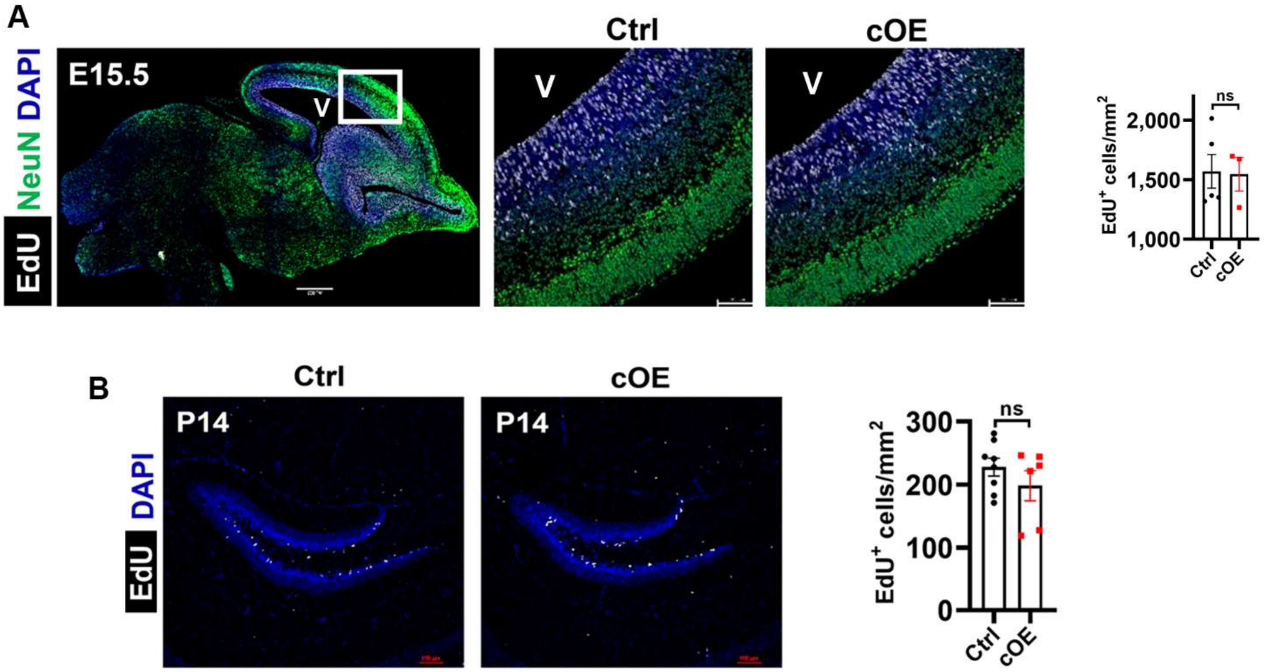


**A**, fluorescence images and quantification of NeuN and EdU (2 hours pulse labeling) in the cerebral cortex of E15.5 brain. N=5 Ctrl, 3 cOE, unpaired Student's t test, $t_{(6)}$ = 0.1145 *P* = 0.9126.

**B**, low-magnification confocal images and of EdU (24 hours pulse labeling) and quantification in the hippocampus of P14 mice. N=8 Ctrl, 6 cOE, unpaired Student's test, $t_{(12)}$ = 1.128 *P* = 0.2814.

Scale bars: A, 500µm (low magnification), 100µm (high magnification), B, 100µm.

**Suppl Fig 6 - Serpina3n expression does not affect neuronal and glial cell development.**

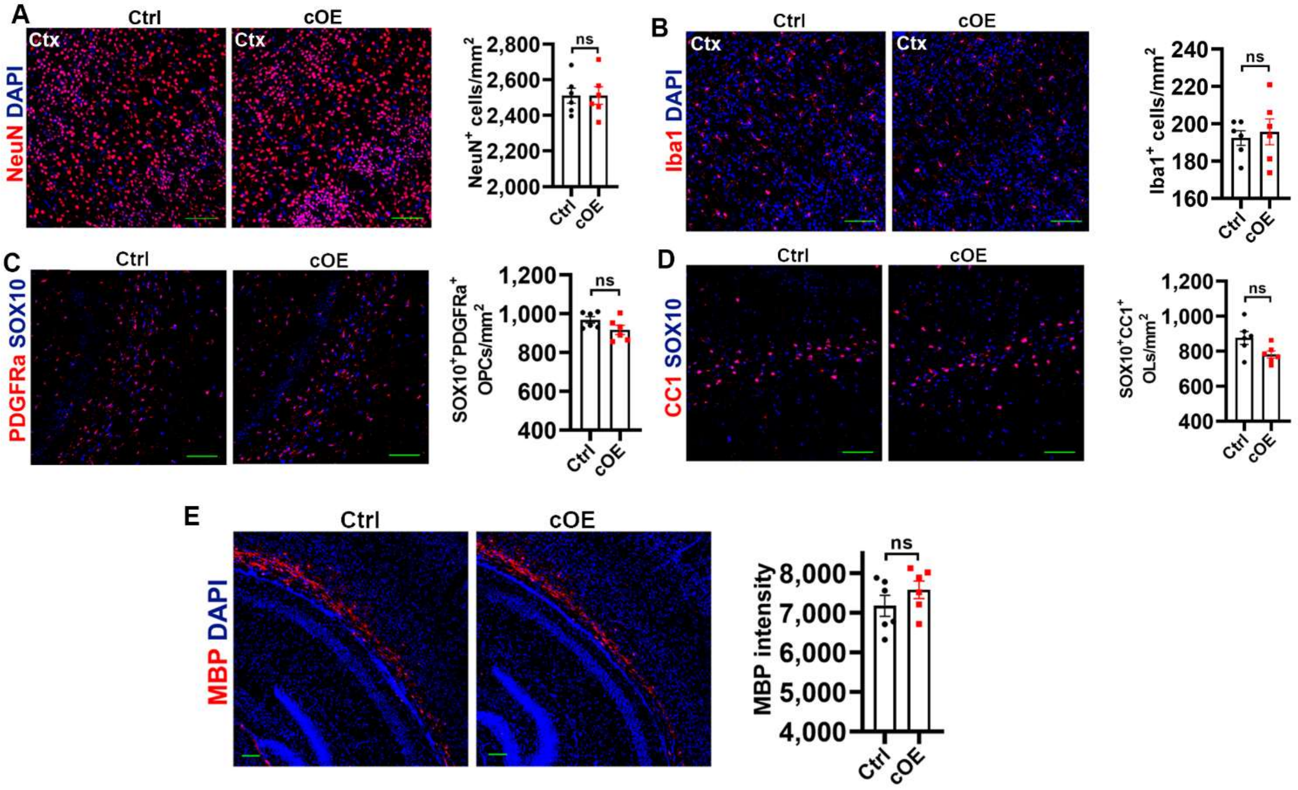


**A**, representative images and quantification of neuronal marker NeuN in the cerebral cortex of P10 mice. N=6, unpaired Student's t test, $t_{(10)}$ = 0.0127 *P* = 0.9901.

**B**, representative images and quantification of microglial marker IBA1 in the cerebral cortex of P10 mice. N=6, unpaired Student's t test, $t_{(10)}$ = 0.4146 *P* = 0.68721.

**C**, representative images and quantification of SOX10 and OPC marker PDGFRa in the subcortical white matter of P10 mice. N=6, unpaired Student's t test, $t_{(10)}$ = 1.807 *P* = 0.1008.

**D**, representative images and quantification of pan-oligodendroglial lineage marker SOX10 and mature oligodendrocyte marker CC1 in the subcortical white matter of P10 mice. N=6, unpaired Student's t test, $t_{(10)}$ = 2.149 *P* = 0.0571.

**E**, representative images and quantification of myelin marker MBP in the cerebral cortex of P10 mice. N=6, unpaired Student's t test, $t_{(10)}$ = 01.176 *P* = 0.2669.

Scale bars=100μm.

**Suppl Fig 7 - Serpina3n expression does not affect BBB integrity during postnatal development**

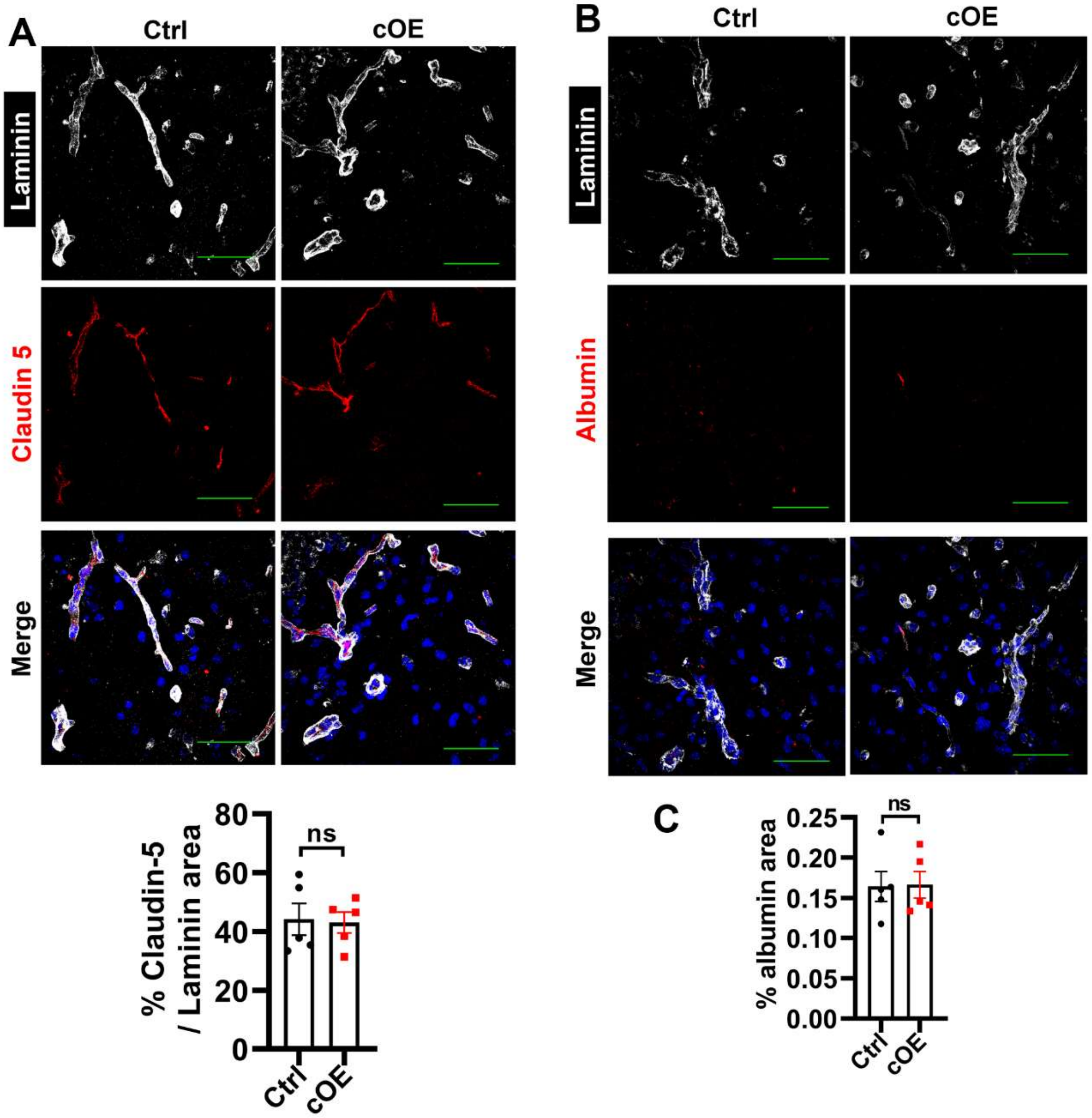


**A**, fluorescence immunostaining of Laminin and tight junction protein Claudin-5 in the hippocampus of P10 mice. N=5, unpaired Student's t test, $t_{(8)} = 0.1702$ $P = 0.8691$.

**B-C**, confocal images and quantification showing absence of Albumin in the hippocampus of P10 mice. N=5, unpaired Student's t test, $t_{(8)} = 0.0865$ $P = 0.9332$.

Scale bars=50μm.

**Suppl Fig 8 - Serpina3n expression does not elicit neuroinflammation or glial activation during homeostasis.**

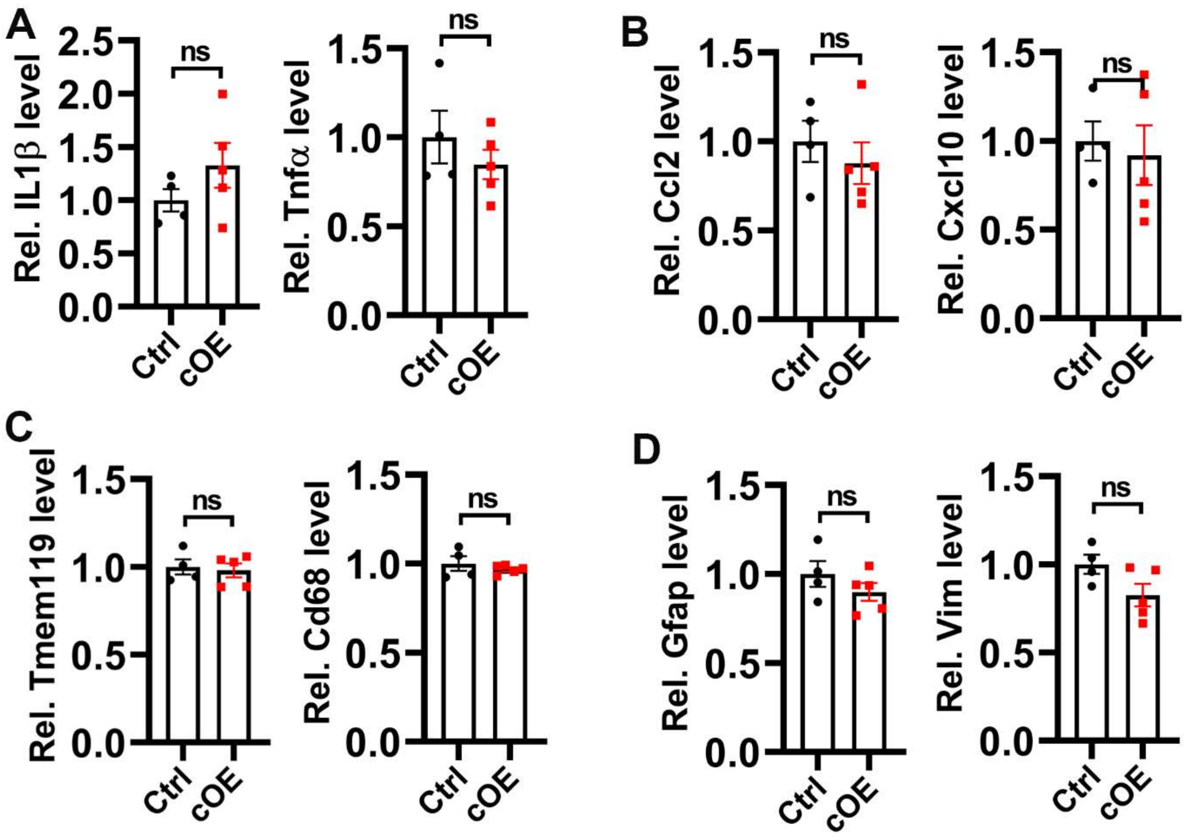


**A**, relative expression of canonical pro-inflammatory cytokine IL1β and Tnfα. Unpaired Student's t test, $t_{(7)} = 1.292$ $P = 0.2375$ IL1β, $t_{(7)} = 0.9592$ $P = 0.3694$ Tnfα,

**B**, relative expression of canonical pro-inflammatory chemokine Ccl2 and Cxcl10. Unpaired Student's t test, $t_{(7)} = 0.7362$ $P = 0.4855$ Ccl2, $t_{(7)} = 0.3755$ $P = 0.7184$ Cxcl10, $t_{(7)} = 0.3408$ $P = 0.7433$ Tmem119, $t_{(7)} = 0.8630$ $P = 0.4167$ Cd68, $t_{(7)} = 1.186$ $P = 0.2742$ Gfap, $t_{(7)} = 2.022$ $P = 0.0827$ Vim.

**C**, relative expression of microglial homeostasis marker Tmem119 and activation marker Cd68. Unpaired Student's t test, $t_{(7)} = 0.3408$ $P = 0.7433$ Tmem119, $t_{(7)} = 0.8630$ $P = 0.4167$ Cd68,

**C**, relative expression of pan-reactive astrocyte marker Gfap and Vim (Vimentin). Unpaired Student's t test, $t_{(7)} = 1.186$ $P = 0.2742$ Gfap, $t_{(7)} = 2.022$ $P = 0.0827$ Vim.

P9 mice (N=4 Ctrl and 5 cOE) were used for RT-qPCR quantification of indicated genes in the brain.

**Suppl Figure 9 - temporal dynamics of SERPINA3N induction in Serpina3n-tdTom reporter mice after neonatal H/I injury**

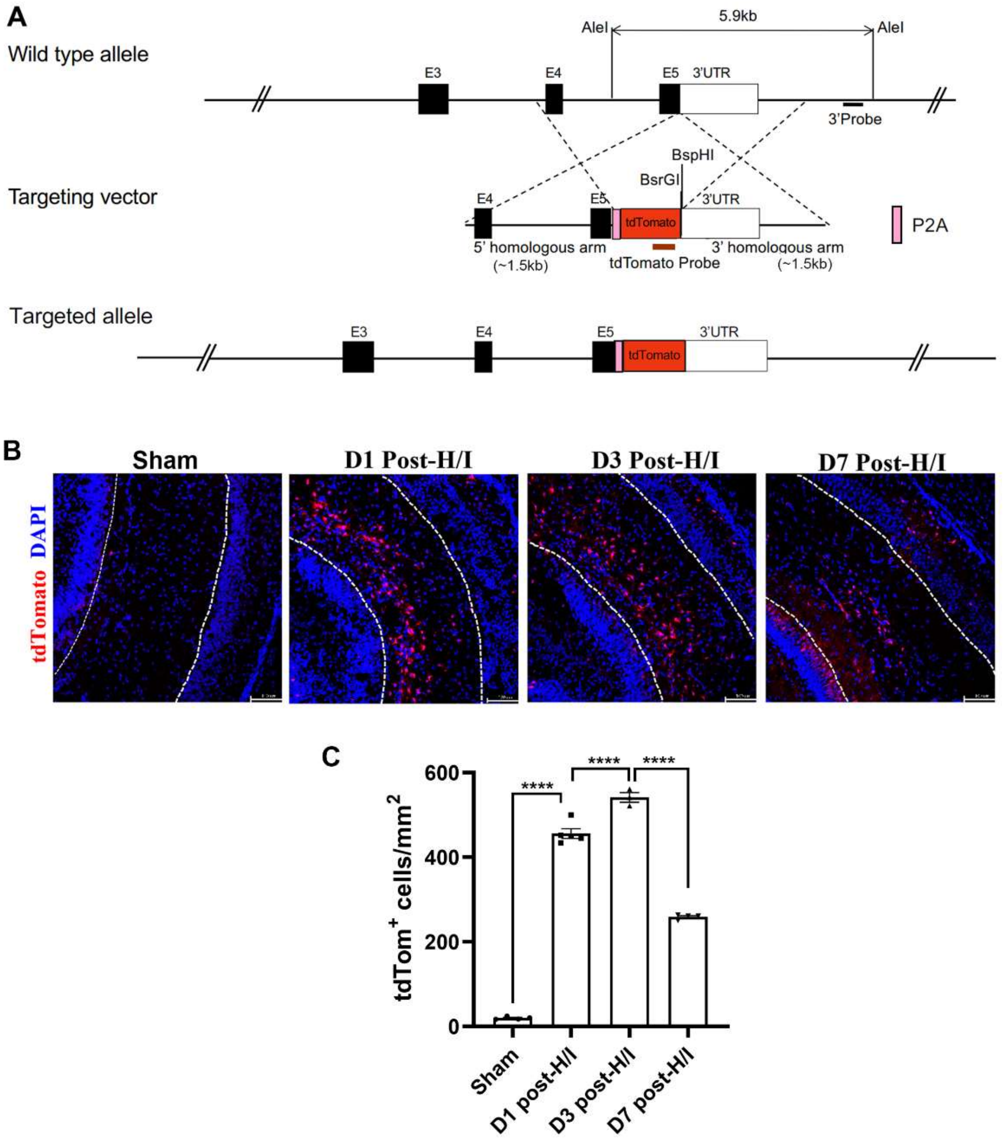


**A**, schematic diagram depicting the constructs of wild type allele of murine Serpina3n, targeting vector, and targeted alleles of tdTomato. The coding sequence of tdTom was knocked into the mouse Serpina3n locus add replaced the stop codon of Serpina3n gene. Therefore, tdTom is expressed only after Serpina3n protein translation. The self-cleavage site P2A between Serpina3n and tdTom ensures the separation of tdTom from endogenous SERPINA3N protein.

**B**, fluorescence immunostaining and quantification showing temporal dynamics of tdTom expression (i.e. SERPINA3N induction) in the hippocampus after hypoxia/ischemia (H/I) injury (Scale bars=100μm). N=3 sham, 5 D1, 3 D3, 4 D7 post-H/I. One-way ANOVA followed by Tukey's multi-comparison test, $F_{(3,12)} = 693.8$ $P < 0.0001$. **** P < 0.0001.
